# TGF/J Regulated Small GTPase RHOV interact with PEAK1 and drive MYC Expression to Promote Cellular Proliferation, Migration and Etoposide resistance

**DOI:** 10.1101/2025.04.18.649622

**Authors:** Annesha Chatterjee, Disha Acharya, Nikita Bhandari, Panchami Bhat, Balkrishna Chaube, Sudhanshu Shukla

**Affiliations:** Cancer Biology lab, Department of Biosciences and Bioengineering, Indian Institute of Technology Dharwad, Dharwad, 580011

**Author notes:** Corresponding author Address for correspondence: Cancer Biology lab, Department of Biosciences and Bioengineering, Indian Institute of Technology Dharwad, Dharwad, 580011.

**Keywords:** Adenocarcinomas, GTPase, RHOV, MYC, TGFβ, Proliferation, Metastasis

## Abstract

1.

Non-small cell lung cancer (NSCLC) remains a leading cause of cancer-related mortality, driven by tumor heterogeneity, metastasis, and therapeutic resistance. While Rho GTPases are well-established regulators of oncogenic processes, the role of the atypical GTPases in NSCLC remains unexplored. Here, we identified RHOV as one of the commonly upregulated Rho GTPases in NSCLC. Analysis of four independent patient cohorts revealed that elevated RHOV expression serves as a robust and independent prognosticator of NSCLC patients specifically early-stage disease. Functionally, RHOV knockdown significantly inhibited cell proliferation, whereas its overexpression enhanced proliferation. Similarly, RHOV depletion suppressed cell migration by disrupting cytoskeletal dynamics, while its overexpression promoted migratory capacity. Mechanistically, we demonstrated that RHOV is a direct transcriptional target of the TGFβ-SMAD3 signaling pathway. RNA-seq analysis identified MYC as a critical downstream mediator of RHOV; RHOV knockdown reduced MYC expression, impairing mitochondrial oxidative phosphorylation and inducing ROS-mediated DNA damage—a phenotype rescued by MYC overexpression. Furthermore, RHOV inhibition sensitized NSCLC cells to etoposide but not doxorubicin. immunoprecipitation coupled with LC-MS revealed PEAK1 as a key interactor of RHOV. The RHOV-PEAK1 complex proved essential for NSCLC proliferation, as PEAK1 silencing abolished RHOV- driven MYC upregulation and tumor growth. This axis sustains MYC levels and activates PI3K/MAPK signaling. Intriguingly, PEAK1 depletion elevated TGF-β levels, which suppressed RHOV expression, establishing a negative feedback loop wherein PEAK1 maintains RHOV by inhibiting TGF-β signaling. Collectively, our findings establish RHOV as a prognostic biomarker and a driver of NSCLC progression via the RHOV-PEAK1-MYC axis, highlighting its potential as a therapeutic target.

**Highlights:**

1. RHOV upregulation predicts poor NSCLC survival, particularly in early-stage disease.
2. The RHOV-PEAK1 interaction is crucial for NSCLC growth and cell migration.
3. RHOV inhibition sensitizes NSCLC cells to Etoposide treatment.
4. RHOV expression is sustained via a PEAK1-TGFβ negative feedback loop.

## 2. Introduction

Lung cancer remains the leading cause of cancer-related mortality worldwide, with non– small cell lung cancer (NSCLC) representing approximately 84% of all cases[1]. Despite significant advancements in targeted therapies and immunotherapy, NSCLC continues to exhibit notable genetic heterogeneity, aggressive metastatic behavior, and frequent development of therapeutic resistance [2]. These challenges highlight the need to discover novel molecular drivers that can act as prognostic markers and therapeutic targets.

Small GTPases of the Rho family are crucial molecular switches that regulate a wide range of cellular processes, including actin cytoskeleton dynamics, cell adhesion, migration, and proliferation [3] . While the roles of classical Rho GTPases such as RHOA, Rac1, and Cdc42 [4] are well established in cancer progression, emerging evidence highlights the importance of atypical Rho GTPases—particularly RHOV—as key modulators of tumorigenesis [5,6]. Unlike classical Rho proteins, RHOV features unique regulatory characteristics, such as non-canonical GDP/GTP cycling and distinct post-translational modifications, indicating specialized functions in oncogenic signalling [7].

Multiple independent studies have shown that RHOV is overexpressed in NSCLC tumours compared to adjacent normal tissues, and its elevated levels are linked to poor patient survival [5,7–9]. Functional analyses in lung cancer cell lines indicate that silencing RHOV significantly reduces cell proliferation and migration [5,6]. Furthermore, RHOV-driven activation of key oncogenic pathways, particularly the Jun N-terminal kinase (JNK)/c-Jun cascade, has been associated with promoting both tumour growth and metastatic dissemination in NSCLC [6]. RHOV signalling plays a crucial role in cellular behaviour and is heavily influenced by its interactions with adaptor proteins. Recent studies on the molecular regulation of RHOV have identified that its N-terminal proline-rich motif is responsible for binding to SH3 domain-containing proteins, including GRB2 and NCK2[10]. These interactions significantly modulate downstream signalling pathways, thereby refining the oncogenic output of RHOV. However, the upstream regulation of RHOV and its interactome and downstream signalling which influences the carcinogenesis processes are not yet deciphered.

Here, we investigated RHOV in NSCLC and found it crucial for tumor proliferation and migration. TGFβ negatively regulated RHOV, and transcriptomic profiling revealed MYC-related genes as the most downregulated in RHOV knockdown samples. MYC downregulation was confirmed, and its overexpression rescued proliferation and metabolic deficits caused by RHOV loss. Mass spectrometry identified PEAK1 as a RHOV interactor, suggesting RHOV modulates MYC via PEAK1. These findings establish RHOV as a prognostic biomarker and the RHOV–PEAK1–MYC axis as a therapeutic target to disrupt NSCLC growth, metabolism, and resistance.

## 3. Materials and Methods

### Expression and Clinical Data

RNA-sequencing (RNA-seq) expression data from The Cancer Genome Atlas (TCGA) project were downloaded from the Broad Institute GDAC Firehose platform (https://gdac.broadinstitute.org). Data pertaining to a University of Michigan cohort were utilized based on a previous study [8]. Gene expression data for an East Asian cohort (EAC) were obtained as described by Chen et al.[11] Additionally, relevant microarray datasets were downloaded from the Kaplan-Meier (KM) Plotter online database (www.kmplot.com).

### Statistical Analysis

To compare expression levels between tumor and normal, the non-parametric Mann-Whitney U-test was employed. Survival analyses, specifically Cox proportional hazards regression modeling, were performed using the survival package within the R statistical environment. Kaplan-Meier survival curves were generated and visualized using GraphPad Prism version 10 (GraphPad Software, La Jolla, CA, USA).

### Plasmids

Primers targeting RHOV were designed using Primer3Plus (https://www.primer3plus.com). Two RHOV silencer predesigned siRNAs were ordered from Ambion ( Cat #AM16708; ID 120714 &120715). The shRNA sequences for RHOV knockdown were generated via the GPP Web Portal (https://portals.broadinstitute.org/gpp/public/) and subsequently cloned into the lentiviral vector pLKO.1 puro (Addgene, 8453). RHOV overexpression construct cloned in pcDNA3.1^+^C-(K)DYK was ordered from GenScript( Product ID- OHu11609D). To study overexpression, we cloned the RHOV overexpression construct in the lentiviral vector pLenti-cMYC-DDK-Puro (Addgene # 123299). Luciferase assay was performed using the reporter construct pGL4-UPRE-luc2P-Hygro (Addgene, #101788).DNA damage responses were studied using Apple 53-BP1trunc (Addgene #69531).

### Cell Culture

The cell lines used in this study were obtained from the American Type Culture Collection (ATCC) and maintained in the recommended growth medium supplemented with 10% fetal bovine serum (FBS) (Gibco #10270106)) and 1% Penicillin-Streptomycin (PenStrep) (Gibco #15140122) at 37°C in a 5% CO[ environment. The Lenti-X 293T cell line (Takara #632180) was utilised for lentiviral production through transfection using the Xfect Transfection Reagent (Takara #631317), following the manufacturer’s protocol. Lentiviral infection was facilitated by polybrene-mediated transduction, and successfully infected cells were selected using puromycin (3 µg/mL).

For functional assays, in a 96-well plate, cell proliferation was monitored by plating 3,000 cells per well and measuring confluency over time using the Incucyte Live-Cell Imaging System. For the wound healing assay, 40,000 cells per well were seeded, and a scratch was introduced using the Incucyte WoundMaker. Colony formation assays were conducted by plating 1,000 cells per well, and after 14 days of incubation, colonies were fixed and stained with 0.25% Crystal Violet for visualisation. Plasmid DNA transfection was done using Lipofectamine 3000 transfection reagent ( Thermo Fisher # L3000008). siRNA transfection was done using Lipofectamine RNAiMAX transfection reagent ( Thermo Fisher #13778075).

### Genomic DNA extraction

Lyse cell sample using 25 μL Proteinase K and up to 200 μL cells. Add 200 μL Lysis Buffer BQ1 to the samples and vortex the mixture vigorously (10 – 20 s). Incubate samples at 70 °C for 10 – 15 minutes. The lysate should become brownish during incubation with Buffer BQ1. Adjust DNA binding conditions by adding 200μL ethanol (96 – 100 %) to each sample and vortex again. To bind DNA apply the samples to the NucleoSpin® Quick Pure Columns placed in a Collection Tube and centrifuge 1 min at 11,000 x g. If the samples are not drawn through the matrix completely, repeat the centrifugation at a higher g-force (up to 15,000 x g). Discard Collection Tube with flow-through. Wash the silica membrane by placing the NucleoSpin® Column into a new Collection Tube (2 mL) and add 350 μL Buffer BQ2. Centrifuge 3 min at 11,000 x g. Discard Collection Tube with flow-through. The drying of the NucleoSpin® Blood Quick Pure Column is performed by 3 min centrifugation. Place the NucleoSpin® Blood Quick Pure Column in a 1.5 mL microcentrifuge tube and add 50μL prewarmed Buffer BE (70 °C). Dispense buffer directly onto the silica membrane. Incubate at room temperature for 1 min. Centrifuge 1 min at 11,000 x g.

### Staining

Phalloidin stain was used to stain actin filaments. To perform phalloidin staining, begin by seeding 1 lakh cells per mL into each well of a 12-well plate containing pre-coated coverslips, dispensing the cell suspension dropwise and shaking in a figure-8 pattern to ensure even distribution. Incubate the cells overnight at 37°C with 5% CO[. The following day, wash each well three times with PBS for 5 minutes each to remove any debris. Fix the cells by adding 500 µL of 4% paraformaldehyde (PFA) to each well and incubating for 30 minutes at room temperature (RT), while protecting the plate from light using aluminium foil. After fixation, aspirate the PFA and wash the cells three times with PBST (PBS + 0.1% Tween-20), each wash for 5 minutes. To permeabilize the cells, add 500 µL of 0.1% Triton X- 100 to each well and incubate for 15 minutes at RT, then aspirate and wash three times with PBST for 5 minutes each. Block non-specific binding by adding 500 µL of 1% BSA in PBS to each well and incubating for 30 minutes at RT. After blocking, wash the wells twice with PBST for 5 minutes each. Prepare a 1X phalloidin stain solution by diluting the stain 1:1000 in the blocking agent, then add 500 µL of this stain to each well and incubate for 90 minutes at RT, again covering with aluminium foil. Following incubation, aspirate the stain and store it at -20°C for reuse. Wash the wells three times with PBST, each for 5 minutes. Clean the slides and coverslips with ethanol, then add 30 µL of mounting dye to a slide and carefully place the coverslips on the mounting medium with the cells facing downward. Incubate the slide for 10 minutes at RT, then seal the edges of the coverslips with colourless nail polish and allow it to dry. Finally, use a fluorescent microscope to visualize the stained cells and examine the actin filaments.

### Promoter cloning

Primers were designed to amplify the RHOV promoter from the genomic DNA. Then the amplicon was cloned into the pMD20_T vector. Then the luciferase reporter vector- pGL4- UPRE-luc2-Hygro was digested using NheI and HindIII. The promoter cloned in the T vector was PCR amplified and compatible restriction sites were added on either side of the amplicon. After amplification, the amplicon was digested again with NheI and HindIII to make the sides compatible for ligation. The insert and vector were ligated using a 1:8 molar ratio using T4 DNA ligase. Then transformed into DH5alpha-competent cells.

### MTT Assay

Cells were plated in 96-well plates, then post 24 hrs of plating, 10ul of MTT was added and cells were incubated at 37 °C for 4 hrs. After that DMSO was added and the absorbance was read, which measures cell viability based on mitochondrial activity.

### Treatments

MG132 (Proteasomal inhibitor, 10uM) was used and cells were collected at an interval of 4hrs and analysed further. TGF β1treatment (10uM) was administered for 24 hrs. and TGF β1 inhibitor (SelleckChem, S2805) (10uM) was administered for 48 hrs. STAT3 inhibitor (10uM) was administered for 24 hrs.

### Luciferase Assay

The cells were transfected with the RHOV promoter cloned luciferase vector, after a 24–48- hour incubation, cells were lysed directly in the plate using Passive Lysis Buffer. Firefly luciferase activity is then measured by adding Luciferase Assay Reagent II, followed by the addition of Stop & Glo® reagent to quench the firefly reaction and initiate Renilla luciferase activity. We assessed RHOV promoter activity using the Dual-Luciferase® Reporter Assay System (Promega #E1910).

### Measuring ATP levels and oxygen consumption rate

We assessed oxidative phosphorylation by quantifying cellular ATP levels using the CellTiter-Glo® Luminescent Cell Viability Assay (Promega #G7570). We measured cellular oxidative phosphorylation by assessing extracellular oxygen consumption using Abcam’s Extracellular Oxygen Consumption Reagent (catalogue #ab197242).

### qRT-PCR

Total RNA was extracted by suspending cells in RNAiso Plus reagent (Takara #9108) and isolating RNA using chloroform as per the manufacturer’s protocol. The purified RNA was quantified and reverse-transcribed into cDNA using the iScript cDNA Synthesis Kit (Bio-Rad #1708841). For qRT-PCR, the synthesized cDNA served as a template in reactions prepared with Universal SYBR Green Mix (Bio-Rad #172571), and TPT primers were included as internal controls. The data analysis was performed using the Bio-Rad CFX Maestro software.

### Co-Immunoprecipitation Assay

Cells grown in T25 flasks were first washed with cold 1X PBS. Next, 300 µL of trypsin was added to detach the cells, which were then collected and centrifuged at 1,000 g for 4 minutes at room temperature. The resulting cell pellet was resuspended in 500 µL of IP lysis buffer, and all subsequent steps were performed on ice. The lysate was gently mixed for 30 minutes on ice, then centrifuged at 13,000 rpm for 15 minutes at 4°C. The supernatant was transferred to a fresh tube, and 50 µL was set aside as input for Western blot analysis. For the antibody-protein binding step, 450 µL of the pre-chilled protein lysate was transferred to a new 1.5 mL tube and mixed with 3 µL of FLAG antibody (Anti-Flag M2, IgG1, Mouse). This mixture was incubated overnight at 4°C with mild rotation. Following antibody binding, protein G beads were prepared in a fresh, pre-chilled 1.5 mL tube placed on a magnetic stand. Forty microliters of Magne Protein G beads (Promega) were added, and the tube was placed in a magnetic field for 30 seconds to pull the beads to the side. After the beads were settled, the storage solution was carefully removed without centrifugation. The beads were then washed with 100 µL of IP lysis/wash buffer, and after a brief magnetic pull (30 seconds), the wash buffer was removed. The beads were resuspended in 100 µL of IP lysis/wash buffer and then added to the tube containing the protein-antibody conjugate. This mixture was incubated for 2 hours at 4°C with mild rotation. For elution, the sample was placed back on the magnetic rack for 30 seconds to fully collect the beads, and the supernatant was discarded. The beads were then washed with cold 1X PBS (after gently mixing by inversion), and the PBS was removed once the beads were magnetically collected. Finally, 35 µL of 2x Laemmli Sample Buffer containing β-mercaptoethanol was added to the beads, and the sample was boiled at 100°C for 3 minutes. Twenty microliters of the eluate were loaded for SDS-PAGE, followed by Western blot confirmation.

### Western Blot

Proteins were extracted by suspending cell pellets in Pierce IP Lysis Buffer (Thermo Fisher #87787) supplemented with 1× Protease Inhibitor Cocktail (Thermo Fisher #87787). Following lysis, the samples were centrifuged at 13,000 × g for 12 minutes at 4°C, and the supernatant containing soluble proteins was collected. Protein concentrations were determined using the BCA Protein Assay Kit(Pierce BCA protein estimation kit #23227). A total of 40 µg of protein from each sample was resolved on a 12% polyacrylamide gel via SDS-PAGE and subsequently transferred to a PVDF membrane (Bio-Rad #1620177) using a wet transfer system. The membrane was blocked with 5% blocking powder (G-Biosciences #786-011) to prevent nonspecific binding and incubated with the appropriate primary antibodies overnight at 4°C.

After thorough washing, the membrane was incubated with the corresponding secondary antibodies for 90 minutes at room temperature. Following additional washes, the membranes were visualised using the FemtoLucent Plus HRP detection system (G-Biosciences #786- 003), and protein bands were imaged for analysis.

### Flow Cytometry

To analyse the cell cycle profile, cells were stained with Krishan Buffer containing propidium iodide (PI)(SRL #25535-16-4) and incubated in the dark for 10 minutes. Stained cells were then analysed using an Attune Flow Cytometer.

### Data Analysis

All statistical analyses were performed using GraphPad Prism. RNA sequencing data were processed and analysed on a Linux platform using R, with differential expression performed using DESeq2 and subsequent Gene Set Enrichment Analysis (GSEA). Mass spectrometry data were processed using MaxQuant for protein identification and quantification, followed by data validation and further analysis with PeptideShaker and R. All experiments were conducted in triplicate, and results are expressed as mean ± standard deviation, with statistical significance defined as p < 0.05.

## 4. Results

### RHOV is overexpressed in many cancers including NSCLC

Small GTPases, critical regulators of cellular signaling, play pivotal roles in cancer progression, yet many remain underexplored [12]. While the Ras is well-characterized for its oncogenic mutations driving proliferation and survival [13], Rho GTPases—a subfamily governing cytoskeletal dynamics, migration, and invasion—are increasingly linked to metastasis but remain understudied. Although key members like RhoA, Rac1, and Cdc42 are implicated in tumor growth and motility [14], lesser-known Rho proteins and other GTPase families may harbor unexplored roles in tumorigenesis, drug resistance, or tumor microenvironment interactions. To systematically investigate Rho GTPase dysregulation in cancer, we analyzed expression data from TCGA and microarray datasets spanning 12 prevalent cancer types and corresponding normal tissues. Comparative profiling of 21 Rho family members revealed genes such as RAC1, RAC3, and CDC42 as commonly overexpressed, while RHOBTB family members were frequently downregulated. Notably, RHOV emerged as one of the most consistently overexpressed genes across cancers, including non-small cell lung cancer (NSCLC) (**Figure 1A**). Given its limited prior investigation in carcinogenesis, we prioritized RHOV for detailed functional exploration to uncover its role in tumor biology.

**Figure 1:**
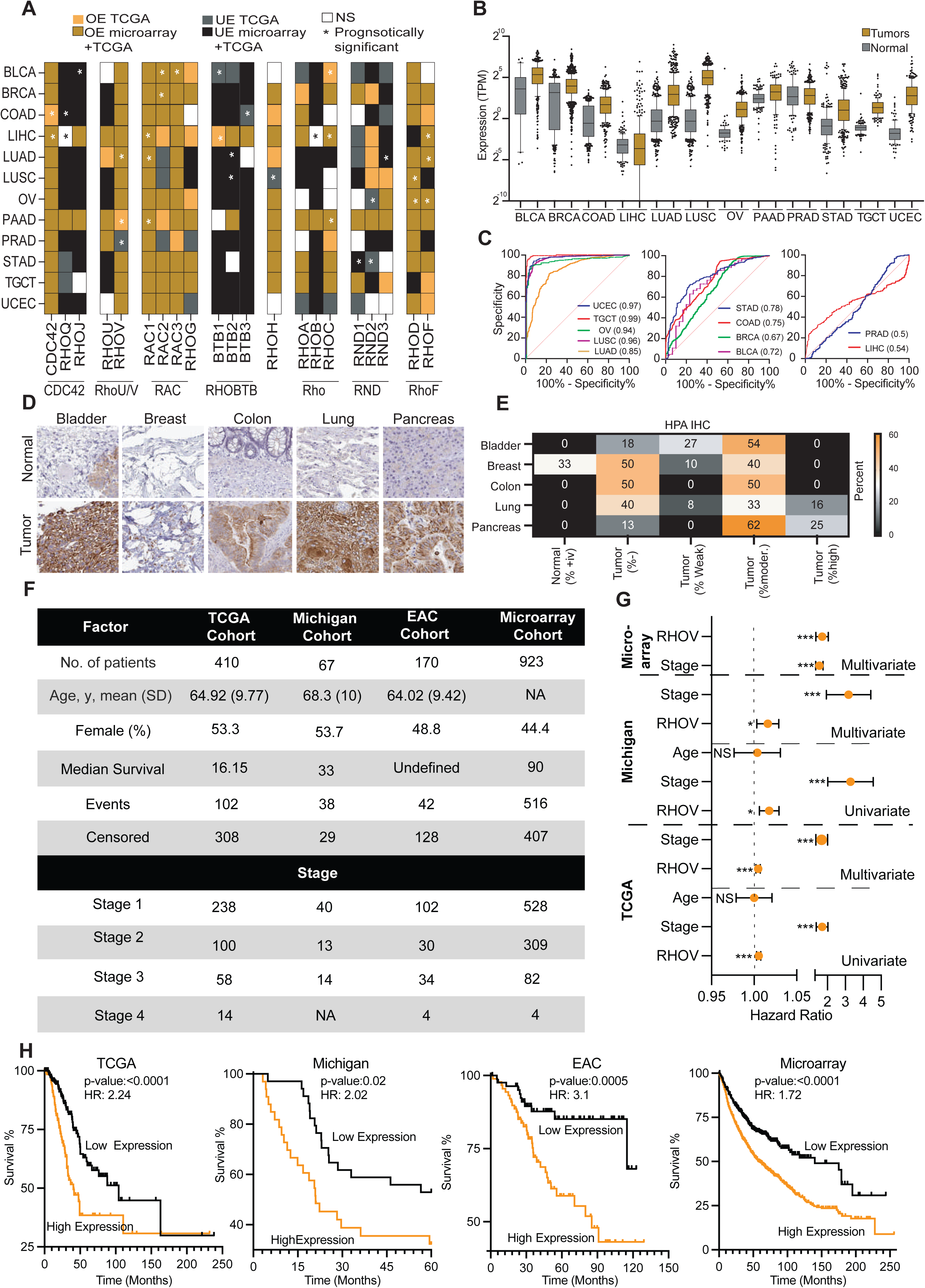
RHOV is overexpressed in multiple solid tumours and correlates with poor prognosis. (A) Heatmap summarizing differential expression of Rho family GTPases across 12 TCGA cancer types and microarray datasets. Overexpression (OE) and under-expression (UE) are denoted in orange and black respectively. Asterisks (*) indicate a statistically significant correlation with prognosis. (B) Boxplot of RHOV transcript levels (TPM) comparing tumor and normal tissues across 12 TCGA cancers. Expression is significantly elevated in tumor tissues of LUAD (P = 2.6 × 10[¹[), LUSC (P = 3.7 × 10[¹³), and PAAD (P = 1.1 × 10[[), Wilcoxon rank-sum test. (C) Receiver Operating Characteristic (ROC) curves showing the diagnostic ability of RHOV expression to distinguish tumors from normal tissues. Area under the curve (AUC) values are highest in UCEC (AUC = 0.97), TGCT (AUC = 0.91), and LUAD (AUC = 0.86), supporting the utility of RHOV as a tumour-specific biomarker. (D) Representative immunohistochemistry (IHC) images from the HPA comparing RHOV protein expression in normal and tumour tissues of the bladder, breast, colon, lung, and pancreas. While normal tissues show absent or weak staining, tumour sections—especially lung and pancreas—exhibit markedly elevated RHOV expression. (E) Quantification of RHOV protein expression in tumours and normal tissues, categorized by staining intensity (Negative, Weak, Moderate, High). (F) Kaplan–Meier survival analyses correlating RHOV expression with overall survival in NSCLC across TCGA, microarray, Michigan and EAC cohorts. In All the data sets, high RHOV expression is associated with significantly reduced survival (TCGA: HR = 2.83; P = 1.0 × 10[[). (Michigan; HR = 3.31; P = 1.0 × 10[[), ( EAC: HR = 3.16; P = 5.0 × 10[[) and (Microarray: HR: 1.72, P <0.0001).

Further explorations of RHOV expression showed that RHOV is significantly overexpressed in many cancer (except prostate adenocarcinoma) (**Figure 1B**). Further validation through Oncomine analysis—utilising multiple datasets supported the conclusion that RHOV is prominently overexpressed in NSCLC (**Supplementary Figure 1A and B**). Furthermore, ROC analysis showed that RHOV has strong specificity and sensitivity in detecting tumour from normal samples for NSCLC (**Figure 1C**). Protein expression data as analysed by immunohistochemistry in cancer samples and corresponding to normal samples also confirmed high expression of RHOV in cancer compared to normal (data obtained from HPA) (**Figure 1D**).

### RHOV is a robust predictor of NSCLC patients survival

The consistent overexpression of RHOV in NSCLC, coupled with its high specificity and sensitivity in distinguishing tumor from normal tissue, suggests its potential as a clinically relevant prognostic biomarker. To explore this, we analyzed four independent cohorts of NSCLC patients, totalling 1540 individuals with detailed clinical data (**clinicopathological features summarized in Figure 1F**). Since the East Asian cohort (EAC) lacked defined median survival, we focused our prognostic analysis on the remaining three cohorts using univariate Cox regression. This analysis revealed RHOV overexpression as a significant predictor of reduced survival in all three cohorts (**Figure 1G**), a finding consistent with the established prognostic significance of advanced tumor stage. Importantly, multivariate Cox regression, incorporating both RHOV expression and tumor stage, demonstrated that RHOV remained an independent predictor of survival (**Figure 1G**). Furthermore, when patients were stratified based on RHOV expression levels, high RHOV expression was consistently associated with significantly poorer survival across all four cohorts (**Figure 1H**), reinforcing its potential as a robust prognostic marker in NSCLC. Prognosis is crucial for guiding therapy decisions in patients with early-stage NSCLC [8]. We investigated the prognostic role of RHOV expression in low-grade NSCLC patients across four datasets. In all datasets, RHOV expression was found to be a significant predictor of prognosis (**Supplementary Figure 1B**), indicating its potential usefulness as a prognostic marker for NSCLC.

### RHOV is required for the NSCLC cell growth

To investigate the functional impact of RHOV overexpression on NSCLC growth, we first assessed RHOV expression across a panel of NSCLC cell lines. Immunoblotting and qRT-PCR revealed detectable RHOV expression in most cell lines, with the exception of H1299 (**Supplementary Figure 2A-B**). To elucidate its oncogenic role, we performed transient and stable RHOV knockdown in lung adenocarcinoma (LUAD) models (H23 and A549 cells). Transient siRNA-mediated silencing achieved >80% reduction in RHOV mRNA and protein levels, resulting in significant suppression of cell proliferation of A549 (**Supplementary Figure 2C-D**) and H23 cells (**Supplementary Figure 2E-F**). Similarly these cells also showed significant reduction in colony formation (**Supplementary Figure 2G-H**). Consistent with these findings, stable RHOV knockdown using shRNA constructs (cloned into the pLKO.1-puro lentiviral vector) markedly impaired proliferation and clonogenicity in both A549 (**Figure 2A-D**) and H23 cells (**Figure 2E-H**).

**Figure 2:**
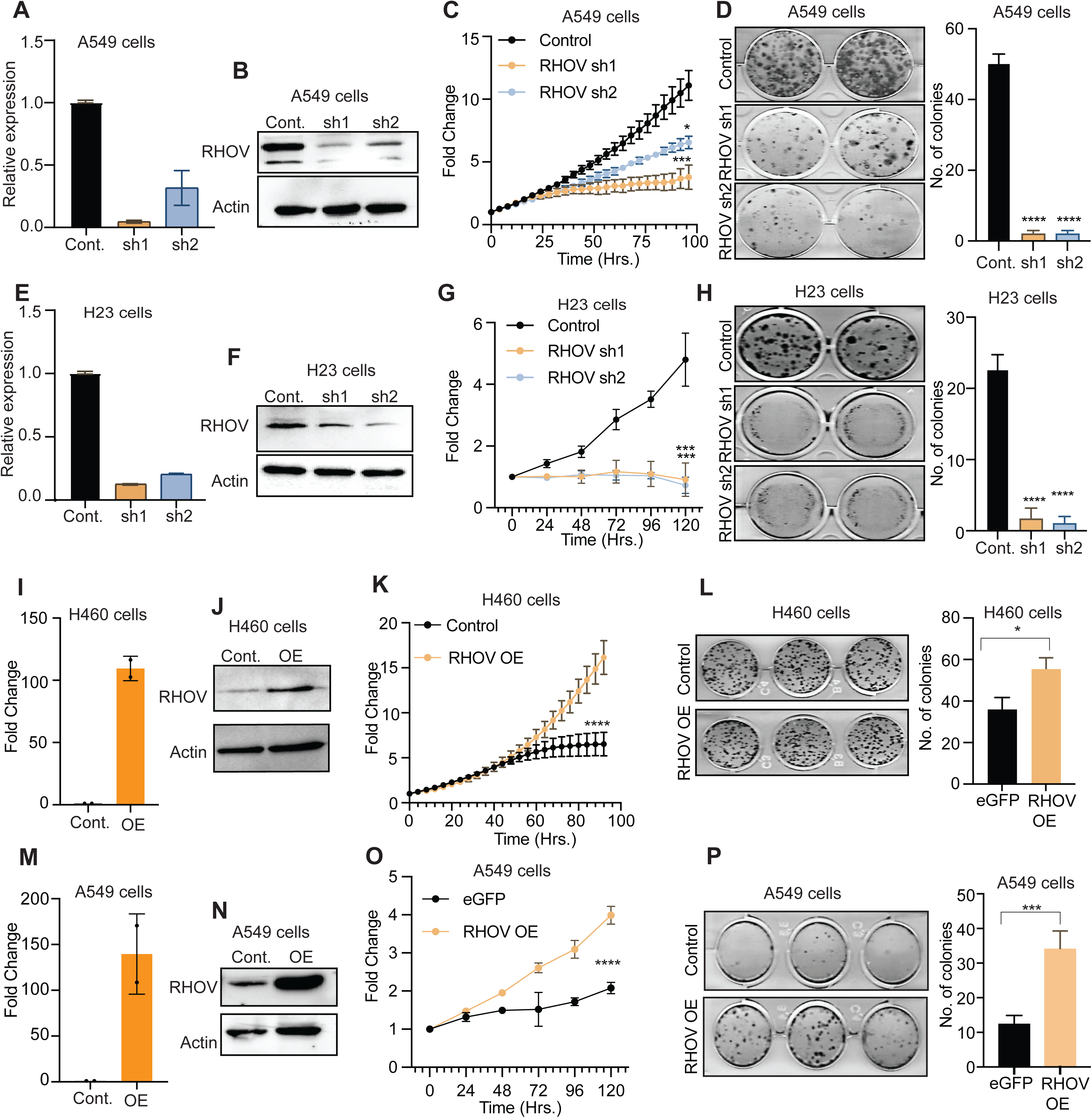
RHOV promotes proliferation and clonogenicity in non-small cell lung cancer (NSCLC) cell lines. (A–D) Knockdown of RHOV in A549 cells using two independent shRNA constructs (sh1, sh2) significantly reduced RHOV mRNA (A) and protein levels (B) as confirmed by qRT-PCR and Western blotting. Actin was used as a loading control. (C) Cell proliferation was assessed by a time-course assay over 96 hours. RHOV knockdown led to significantly reduced growth compared to control cells (Two-way ANOVA with Sidak’s test; ****P < 0.0001). (D) Colony formation assay shows a reduction in clonogenic capacity in RHOV-depleted cells; quantification shows near-complete loss of colony-forming ability (****P < 0.0001, one-way ANOVA). (E–H) RHOV knockdown in H23 NSCLC cells similarly led to a robust reduction in RHOV expression (E–F) and significantly impaired proliferation (G, ****P < 0.0001) and colony formation (H, ****P < 0.0001), confirming the essential role of RHOV in NSCLC cell growth across cell lines. (I–L) Stable overexpression of RHOV in H460 cells resulted in a ∼100-fold increase in RHOV mRNA expression (I) and marked protein overexpression as confirmed by immunoblotting (J). (K) RHOV overexpression significantly enhanced proliferation over 96 hours (****P < 0.0001). (L) RHOV-OE cells formed significantly more and larger colonies compared to control (eGFP- expressing) cells (*P < 0.05, unpaired t-test). (M–P) Similar phenotypes were observed in A549 cells overexpressing RHOV. qPCR (M) and Western blotting (N) validated RHOV overexpression. (O) RHOV-OE cells proliferated significantly faster than controls (***P < 0.001, two-way ANOVA), and (P) formed significantly more colonies in clonogenic assays (***P < 0.001, unpaired t-test).

Conversely, RHOV overexpression in H460 cells (which exhibit moderate endogenous RHOV levels) enhanced oncogenic phenotypes. Ectopic RHOV expression (**Figure 2A** and **B**) significantly accelerated proliferation (**Figure 2C**) and increased colony formation (**Figure 2D**). RHOV overexpression showed similar results in A549 cells as well (**Figure 2E-H**) confirming its role as a driver of NSCLC growth.

### RHOV Regulates Cancer Cell Migration Through Actin-Dependent Mechanisms

Given that RHOV belongs to the RHO GTPase family—a class of proteins known to regulate cytoskeletal dynamics and cell motility—we investigated its potential role in cancer cell migration. To assess this, we generated RHOV-knockdown systems in A549 and H23 lung cancer cells and performed scratch assays to evaluate migratory capacity. In A549 cells, RHOV knockdown significantly delayed wound closure compared to controls (**Figure 3A, B**), a phenotype corroborated in H23 cells (**Supplementary Figure 3A, B**). Also, RHOV knockdown cells showed decreased cell migration in transwell migration assay (**Figure 3C**).

**Figure 3:**
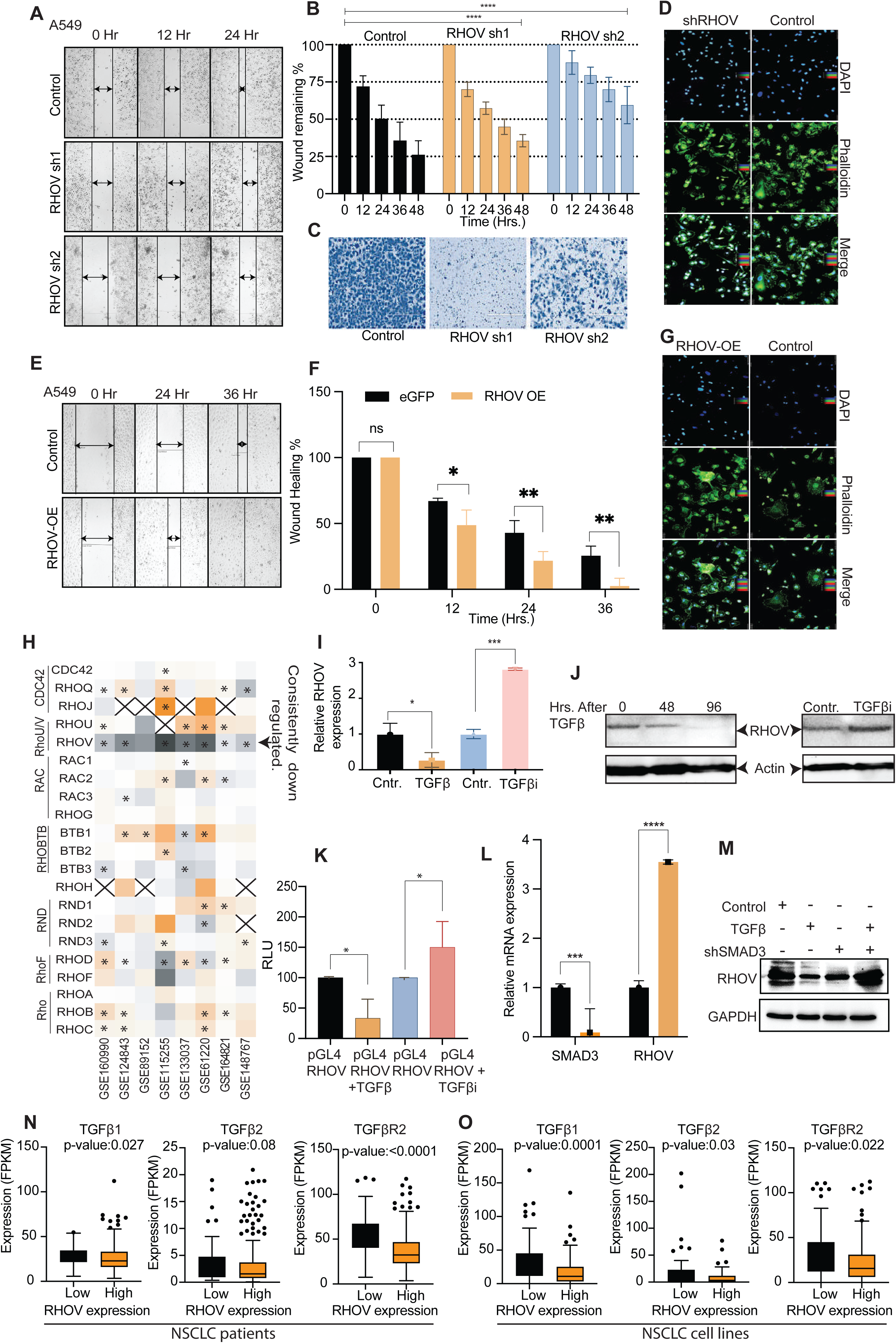
RHOV regulates cell migration via cytoskeletal remodelling and is transcriptionally induced by TGF-**β**–SMAD3 signalling. (A–B) Wound healing assay was performed in A549 cells with stable RHOV knockdown using two independent shRNA constructs (sh1 and sh2). (A) Representative images captured at 0, 12, and 24 hours show delayed wound closure in RHOV-silenced cells. (B) Quantification of wound closure over 48 hours shows significant impairment in migration upon RHOV knockdown (****P < 0.0001, two-way ANOVA). (C) Control cells and shRHOV cells were plated in seeded into the upper inserts in serum-free medium. Migrated cells on the underside of the membrane are then fixed, stained with Crystal Violet. The number of migratory cells were less in shRHOV compared to the control cells. (D) Phalloidin staining in A549 cells reveals disrupted actin filament organization in RHOV- silenced cells, with reduced formation of lamellipodia and filopodia, compared to control. DAPI stains nuclei. (E–F) Wound healing assay in A549 cells stably overexpressing RHOV (OE) compared to eGFP control. (D) Representative images at 0, 24, and 36 hours show accelerated wound closure in RHOV-OE cells. (E) Quantification of wound healing shows significantly enhanced migration in RHOV-overexpressing cells (*P < 0.05, **P < 0.01, two-way ANOVA). (G) Phalloidin staining of RHOV-OE cells reveals enhanced actin polymerisation and increased formation of protrusive structures, including lamellipodia and filopodia, compared to control cells. (H) Heatmap of microarray datasets from publicly available GEO datasets (GSE series) showing RHOV upregulation across multiple contexts, often alongside other Rho family GTPases. RHOV consistently clusters with EMT-associated signatures, suggestive of a role in motility regulation. (I) RHOV mRNA levels were significantly upregulated upon TGF-β1 treatment and significantly downregulated upon treatment with a TGF-β receptor I inhibitor (TGFβi) in A549 cells (*P < 0.05, ***P < 0.001, one-way ANOVA). (J) Western blot analysis shows time-dependent upregulation of RHOV protein upon TGF-β treatment (left), and suppression of RHOV protein by TGFβi (right), confirming transcriptional regulation by the TGF-β pathway. (K) Luciferase reporter assay using the RHOV promoter construct in A549 cells confirms that RHOV promoter activity is significantly increased by TGFβ treatment and attenuated by TGFβi (*P < 0.05, one-way ANOVA). (L–M) RHOV is transcriptionally regulated by SMAD3. (K) qRT-PCR analysis shows RHOV and SMAD3 mRNA levels upon SMAD3 silencing (***P < 0.001, ****P < 0.0001, unpaired t- test). (L) Western blot analysis confirms that shRNA-mediated knockdown of SMAD3 abrogates TGFβ–induced RHOV protein upregulation. (N) Expression of TGBβ1, TGFβ2 and TGFβR2 in TCGA-NSCLC patients divided into two groups based on low and high expression of RHOV. High expression of RHOV was associated with lower expression of TGBβ1, TGFβ2 and TGFβR2 genes. (N) Expression of TGBβ1, TGFβ2 and TGFβR2 in CCLE-NSCLC cell lines divided into two groups based on low and high expression of RHOV. High expression of RHOV was associated with lower expression of TGBβ1, TGFβ2 and TGFβR2 genes.

Cell migration is driven by actin polymerization-dependent membrane protrusion at the leading edge [15]. To determine whether RHOV depletion disrupts this process, we analyzed actin filament organization using phalloidin staining. Consistent with the migration defect, RHOV- knockdown cells exhibited a stark reduction in actin filament-associated protrusive structures compared to controls (**Figure 3D**).

To further validate RHOV’s functional role, we overexpressed RHOV in A549 cells. In contrast to the knockdown phenotype, RHOV-overexpressing cells closed scratch wounds significantly faster than controls (**Figure 3E, F**). Mirroring this enhanced migration, phalloidin staining revealed a pronounced increase in actin-rich membrane protrusions in RHOV-overexpressing cells (**Figure 3G**). Similar results were shown in H460-RHOV overexpressing cells (**Supplementary Figure 3C and D**).

### TGF**β** negatively regulates RHOV through the SMAD3-dependent pathway

We showed that RHOV, a member of the RHO GTPase family, is significantly upregulated in non-small cell lung cancer (NSCLC) and promotes tumor aggressiveness by enhancing cell proliferation and migration. Mechanistically, RHOV facilitates actin filament-dependent membrane protrusions, a critical step in metastatic dissemination. Given that TGFβ is a master regulator of cytoskeletal remodelling and cell motility in cancer [16], we hypothesized that TGFβ signalling may govern RHOV expression or activity to orchestrate migration.

To investigate whether TGFβ regulates RHOV expression, we analyzed eight GEO datasets measuring gene expression after TGFβ treatment. Among the RHO family members, RHOV was uniquely and consistently downregulated across all datasets (**Figure 3H**), a trend further confirmed in five additional datasets (**Supplementary Figure 3E**).

To validate these findings, we treated A549 cells with TGFβ and observed significant repression of RHOV at both mRNA and protein levels (**Figure 3I, J**). Conversely, TGFβ inhibition upregulated RHOV expression (**Figure 3I, J**). To determine if RHOV is a direct transcriptional target of TGFβ signalling, we generated a luciferase reporter construct driven by the RHOV promoter. TGFβ treatment reduced luciferase activity, while TGFβ inhibition increased it (**Figure 3K**), supporting direct transcriptional regulation.

Since SMAD3 is a central mediator of TGFβ signalling [17,18], we examined its role in RHOV regulation. SMAD3 knockdown in A549 cells significantly increased RHOV transcript levels (**Figure 3L**). Moreover, TGFβ failed to suppress RHOV expression in SMAD3-depleted cells (**Figure 3M**), indicating that TGFβ-mediated repression of RHOV depends on the canonical SMAD3 pathway. Furthermore, we checked expression level of TGFβ1, TGFβ2, TGFβR2 in NSCLC patients samples. The expression of these genes were significantly lower in RHOV high expressing samples compared to RHOV low expressing samples (**Figure 3M**). This observation was further confirmed in NSCCL cell lines (**Figure 3N**). These observations confirm that RHOV is direct target of TGFβ pathway.

### RHOV Knockdown Reveals MYC-Mediated Cellular Growth Regulation

To elucidate the mechanism of RHOV-mediated cell growth, we performed RNA sequencing-based expression analysis. The analysis identified a substantial number of differentially expressed genes, with specific counts of downregulated and upregulated genes (**Figure 4A**). Gene Ontology (GO) analysis revealed that RHOV knockdown cells exhibited activation of the DNA damage response among upregulated genes, while downregulated genes were associated with translation and cell cycle processes (**Figure 4B**).

**Figure 4:**
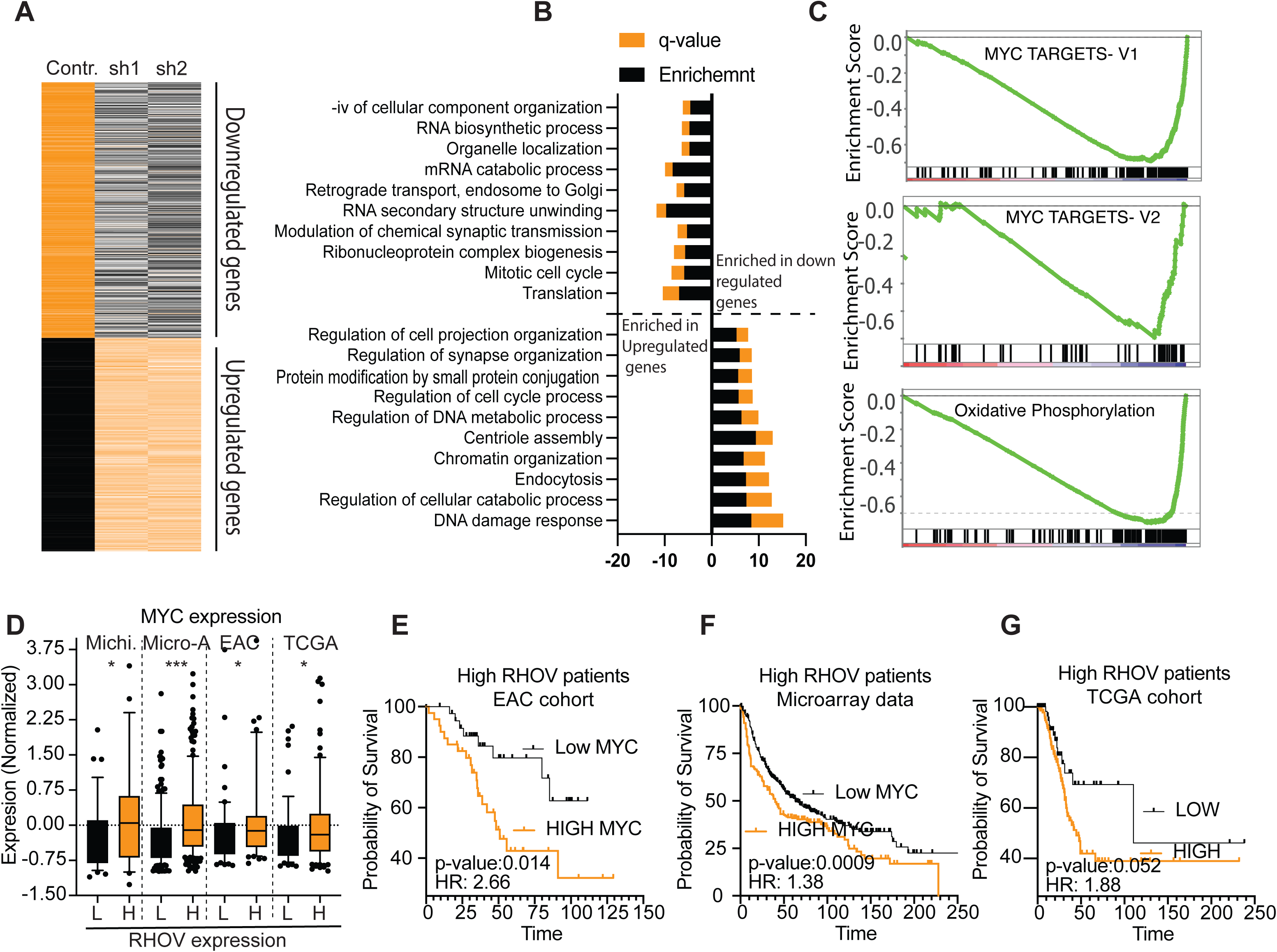
RHOV regulates MYC pathway. (A) Heatmap of differentially expressed genes from RNA-sequencing analysis of A549 cells with stable RHOV knockdown (sh1, sh2) compared to control. Consistent transcriptional alterations across both shRNAs are observed, with enrichment of MYC-related gene signatures among downregulated transcripts. (B) Gene ontology (GO) of differentially expressed genes shows significant downregulation of pathways associated with RNA biosynthesis, mRNA catabolism, ribosomal assembly, and mitotic cell cycle—all known MYC-driven processes. Upregulated pathways include DNA damage response, chromatin organization, and endocytic trafficking. Enrichment scores are represented along with q-values. (C) Gene Set Enrichment Analysis (GSEA) using Hallmark gene sets confirms strong negative enrichment of hallmark MYC target gene sets (V1 and V2) and oxidative phosphorylation genes in RHOV knockdown cells, indicating suppression of both proliferative and metabolic MYC programs. (D) Expression of MYC was compared between high and Low RHOV expressing patients in four NSCLC patients’ datasets. The expression of MYC was significantly higher in high RHOV patients (Mann-Whitney test). (E, F and G) The patients’ samples with high expression of RHOV (which show poor survival, Figure 1H) from three cohort, (E) EAC, (F) Microarray and (G) TCGA were divided into two groups based on MYC expression. The analysis shows that RHOV high patients with low MYC expression survive significantly better than the RHOV high patients with high MYC expression. The p-values and Hazard ration are shown.

Gene Set Enrichment Analysis (GSEA) following RHOV knockdown revealed significant negative enrichment for both MYC target genes and oxidative phosphorylation pathways (**Figure 4C**), suggesting MYC involvement in RHOV-mediated cell growth regulation. To validate this in a clinical context, we analyzed MYC expression across four NSCLC patient datasets. Patients with high RHOV expression exhibited significantly higher MYC RNA levels compared to those with low RHOV expression (**Figure 4D**). Furthermore, within the high-

RHOV patient group (previously associated with poor survival, Figure 1H), those with low MYC expression showed significantly improved survival across three patient cohorts (**Figures 4E, F and G**). These findings suggest that MYC may mediate, at least in part, the observed effects of RHOV in NSCLC patients.

Consequently, we investigated MYC expression levels in RHOV knockdown cells, confirming a significant decrease in MYC expression at both RNA and protein levels (**Figure 5A, B**). To determine whether the reduction in MYC expression was responsible for RHOV-mediated cell growth regulation, we overexpressed MYC in RHOV knockdown cells. Notably, MYC overexpression significantly rescued cell proliferation (**Figure 5C, D**), indicating that MYC is crucial for RHOV-mediated cell growth.

**Figure 5:**
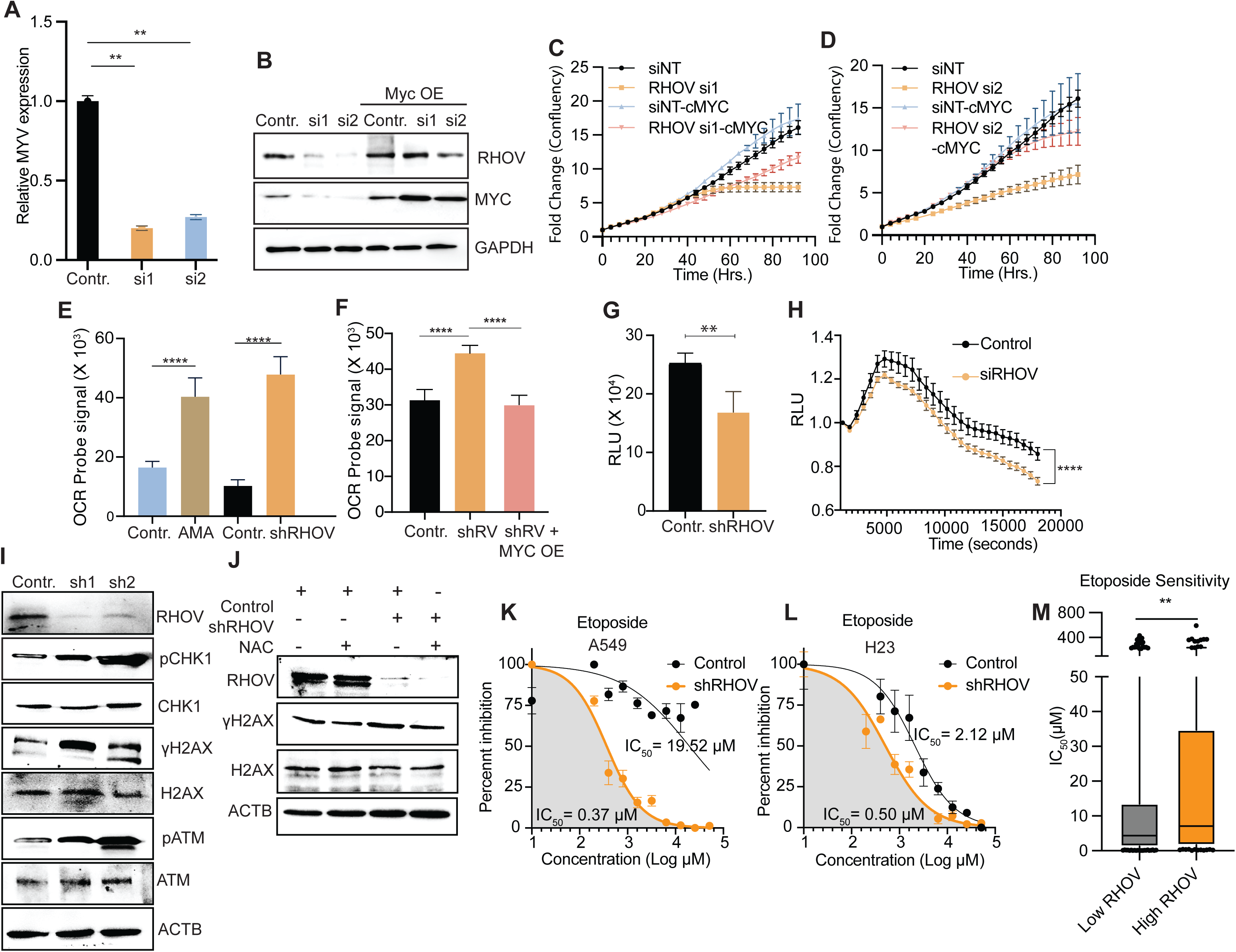
RHOV regulates MYC-dependent transcriptional and metabolic programs in NSCLC. (A–B) qRT-PCR (A) and immunoblotting (B) demonstrate that RHOV silencing (siRHOV-1, siRHOV-2) leads to a significant decrease in MYC expression at both mRNA and protein levels (P < 0.01, one-way ANOVA). Conversely, MYC overexpression rescues protein levels despite RHOV knockdown. (C–D) Cell proliferation assays show that RHOV knockdown impairs proliferation, which is rescued by MYC overexpression in siRHOV-1 and siRHOV-2 backgrounds, confirming MYC as a key downstream effector of RHOV’s pro-growth function (Two-way ANOVA, ****P < 0.0001). (E) Oxygen Consumption Rate (OCR) assay shows that both Antimycin A (AMA, positive control) and RHOV knockdown reduce mitochondrial respiration, indicating suppressed oxidative phosphorylation (****P < 0.0001). (F) OCR rescue upon MYC overexpression is observed in RHOV knockdown cells, confirming MYC’s ability to restore OXPHOS activity (****P < 0.0001, one-way ANOVA). (G) ATP levels measured via CellTiter-Glo 2.0 show significantly reduced luminescence in RHOV-depleted cells (P < 0.01), consistent with impaired mitochondrial ATP production. (H) Real-time ATP kinetics show that RHOV knockdown causes a sustained decline in ATP levels compared to controls (****P < 0.0001, repeated-measures ANOVA), reinforcing the role of RHOV in regulating cellular bioenergetics through MYC. (I) RHOV knockdown induces mitochondrial dysfunction via MYC inhibition, which causes ROS induction (Supplementary figure). DNA damage is one of the common effects of ROS increase. Western blot of RHOV knockdown cells shows increased level of phosphoCHK1, phosphoH2AX and phosphoATM, suggesting increased DNA damage. (J) NAC is a strong quencher of ROS. RHOV knockdown cells, which have high DNA damage show reduction in phosphoH2AX level, suggesting the role of ROS in RHOV induced DNA damage. (K, L) Etoposide induces cytotoxic effect by inhibiting Topoisomerase II and thus DNA repair. The RHOV knockdown cells show significant increase in etoposide sensitivity in both (K) A549 and (L) H23 cells. (M) The etoposide sensitivity of cells lines with high and low expression of RHOV was compared from Genomics of Drug Sensitivity in Cancer database. RHOV high expressing cells showed significant higher IC_50_ than RHOV low expressing cells (p-value= 0.005, Mann- Whitney test).

Given the observed negative enrichment of oxidative phosphorylation genes in RHOV knockdown cells, we further explored mitochondrial metabolism. MYC, a pivotal transcription factor, plays a central role in regulating mitochondrial metabolism by modulating the expression of genes involved in electron transport chain complexes (ETC) [19,20], thereby enabling cancer cells to increase mitochondrial respiratory capacity and ATP generation.

We examined the expression of nuclear-encoded ETC proteins in RHOV knockdown cells and found that the expression of Cytochrome C, DHODH, MCL1, and COX5B was decreased (**Supplementary Figure 4A**). Next, we confirmed reduced mitochondrial activity in RHOV knockdown cells by measuring the oxygen consumption rate (OCR). Compared to control cells, RHOV knockdown cells demonstrated a significant decrease in OCR (**Figure 5E**). Subsequently, MYC overexpression in RHOV knockdown cells resulted in a significant increase in OCR (**Figure 5F**). Consistent with these findings, RHOV knockdown cells exhibited decreased ATP synthesis (**Figure 5G** and **H**). Reduced OCR due to mitochondrial dysfunction often leads to increased ROS production [21]. Therefore, we checked the ROS level in RHOV knockdown cells and observed increased total and mitochondrial ROS production in these cells (**Supplementary Figure 4B, C). I**ncreased ROS is associated with increased DNA damage, and we had observed in pathway analysis that the DNA damage response was the most enriched pathway (**Figure 4B**). Therefore, we checked the expression of various DNA damage repair pathway genes and found increased levels of phospho-CHK1, phospho-H2AX, and phospho-ATM, suggesting increased DNA damage in RHOV knockdown cells (**Figure 5I**). Furthermore, quenching ROS using NAC resulted in reduced phospho-H2AX levels in RHOV knockdown cells, suggesting that MYC-induced ROS is responsible for the increased DNA damage (**Figure 5J**). Since RHOV knockdown cells showed increased ROS-mediated DNA damage, we hypothesized that RHOV knockdown might also make cells more sensitive to chemotherapeutic drugs that induce DNA damage. We checked the effect of etoposide and doxorubicin in RHOV knockdown cells. Interestingly, we found that RHOV knockdown cells drastically increased the etoposide sensitivity of A549 and H23 cells; however, there was no significant change in doxorubicin sensitivity (**Figure 5K** and **L** **and Supplementary Figure 4D**). This observation was further confirmed when we compared the IC_50_ values of etoposide and doxorubicin in cells with high and low RHOV expression using data from the Genomics of Drug Sensitivity in Cancer project. Cells expressing high levels of RHOV were significantly more resistant to etoposide compared to cells expressing low levels (**Figure 5M**). However, there was no difference in doxorubicin sensitivity (**Supplementary Figure 4E**). This could be because etoposide functions by inhibiting Topoisomerase II, which is crucial for repair of ROS induced DNA damage [22]. Inhibition of Topoisomerase II leads to the accumulation of DNA damage in RHOV knockdown cells and exhibiting a synergistic effect. However, doxorubicin induces DNA damage by intercalating into DNA; perhaps because DNA damage levels are already elevated in RHOV knockdown cells, this mechanism does not result in a significant change in their sensitivity to the drug.

### RHOV interacts with PEAK1 to regulate MYC

Building on the evidence that RHOV is crucial for NSCLC cell growth, we sought to characterize its functional interactome. Given RHOV’s atypical role among Rho GTPases in cytoskeletal dynamics and its downregulation by TGFβ, we hypothesized that RHOV interacts with key regulators of cell motility, adhesion, or survival pathways in NSCLC. To elucidate RHOV’s protein interaction network, we employed a FLAG-tagged immunoprecipitation (IP) approach followed by mass spectrometry (MS) analysis (**Figure 6A**). Our network analysis identified key interactors, including PAK1, PAK2, IQGAP1, WASL, and GIT2—proteins involved in cytoskeletal organization, signalling, and motility (**Figure 6B**). The presence of PAK family members suggests a role in pathways governing cell shape and movement, while IQGAP1 and WASL associations highlight RHOV’s involvement in actin cytoskeleton remodelling and cell migration. Further enrichment analysis revealed significant engagement of these interactors in multiple signalling pathways, including ErbB, T cell receptor, Hippo, and MAPK pathways, as well as actin cytoskeleton regulation (**Supplementary figure 5 A, B**). Notably, among the top interactors identified was PEAK1. PEAK1 is a pseudokinase frequently overexpressed in cancers, where it acts as a key signaling molecule promoting cell proliferation, migration, invasion, survival, and therapy resistance [23,24].

**Figure 6:**
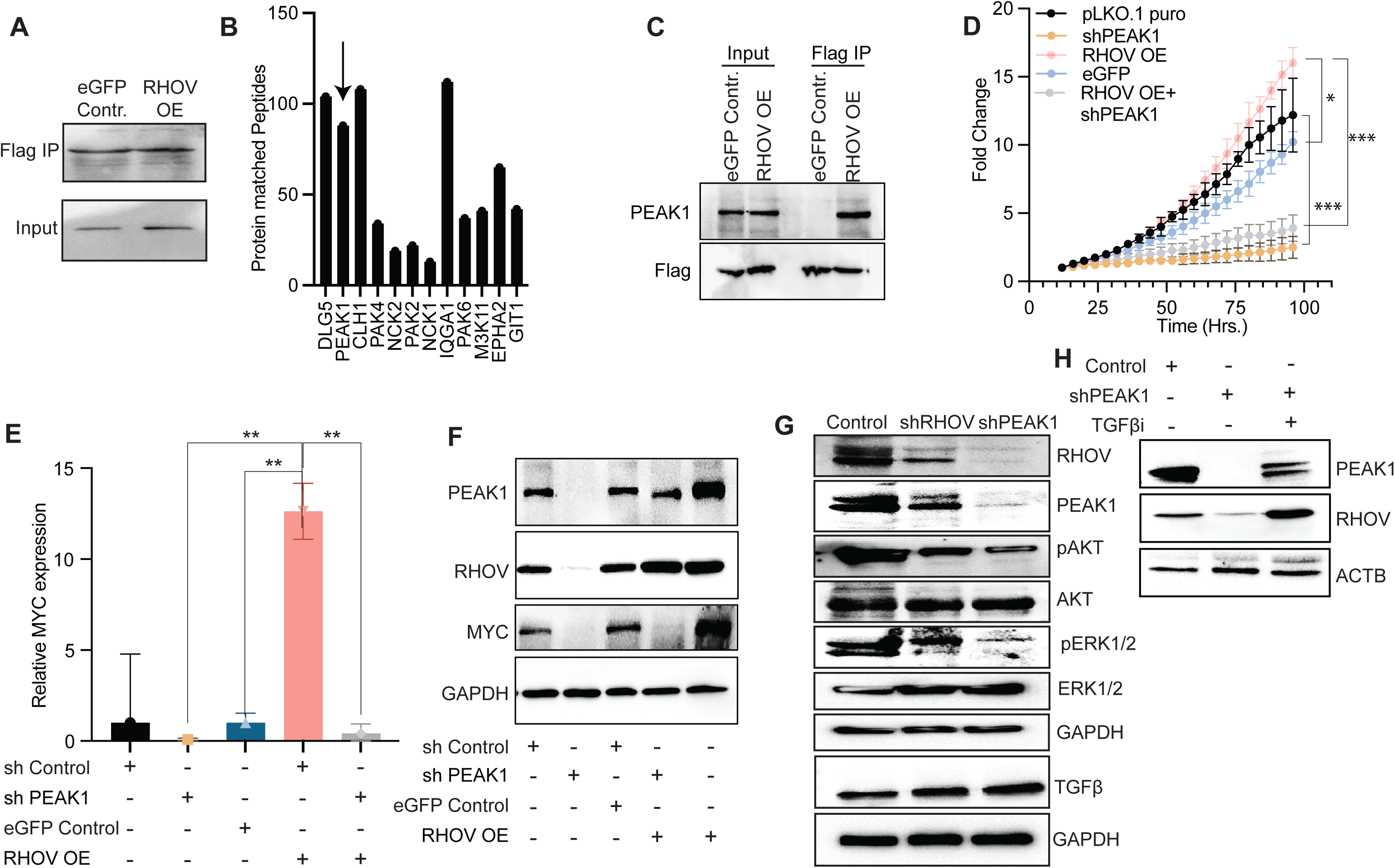
RHOV physically interacts with PEAK1 and cooperatively regulates MYC expression and proliferative signalling. (A) To identify novel RHOV-interacting proteins, we performed co-immunoprecipitation using FLAG-tagged RHOV in HEK293T cells followed by immunoblotting. Efficient immunoprecipitation of RHOV was observed only in the FLAG–RHOV–transfected group, confirming successful pulldown. (B) Mass spectrometry analysis of RHOV immunoprecipitates revealed specific enrichment of multiple known and candidate RHOV interactors. Among the top hits were scaffold protein PEAK1, polarity regulator DLG5, endocytic components CLH1, and PAK family kinases (PAK1, PAK2, PAK4). Previously reported RHOV interactors such as NCK1, NCK2, and IQGAP1 also scored prominently, validating the specificity of the pulldown. (C) Co-immunoprecipitation in RHOV-overexpressing cells validated a specific interaction between PEAK1 and FLAG-RHOV, but not with eGFP control. PEAK1 was detected in the FLAG-IP fraction only in RHOV-OE lysates, confirming a direct or complex-mediated interaction. (D) To determine the functional relevance of this interaction, we assessed proliferation in five experimental conditions: vector control (pLKO.1), PEAK1 knockdown (shPEAK1), RHOV overexpression (RHOV OE), eGFP control, and RHOV OE with PEAK1 knockdown (RHOV OE + shPEAK1). While RHOV overexpression enhanced proliferation, this effect was abolished in the absence of PEAK1, demonstrating that PEAK1 is essential for RHOV-driven proliferation (P < 0.05, **P < 0.001; two-way ANOVA). (E) Quantitative RT-PCR analysis of MYC mRNA levels under the same conditions revealed that PEAK1 knockdown significantly suppressed MYC expression, and abrogated the MYC upregulation seen with RHOV overexpression (P < 0.01, one-way ANOVA), indicating that PEAK1 is necessary for RHOV-mediated activation of MYC. (F) Western blot analysis corroborated these findings at the protein level. MYC expression was downregulated in both RHOV- and PEAK1-depleted cells, while RHOV overexpression increased MYC levels. Critically, PEAK1 knockdown prevented the MYC upregulation induced by RHOV, despite RHOV being overexpressed. (G) To further dissect the downstream signalling consequences, we analyzed the expression and activation status of key oncogenic effectors. Both RHOV and PEAK1 knockdown reduced phosphorylation of AKT (pAKT) and ERK1/2 (pERK1/2), indicating compromised MAPK and PI3K pathway signalling. TGF-β1 protein levels were notably increased upon loss of either RHOV or PEAK1, suggesting that the RHOV–PEAK1 axis negatively regulates TGF-β1 expression, establishing a potential reciprocal regulatory loop. These findings identify PEAK1 as a critical RHOV effector and establish a RHOV–PEAK1–MYC signalling axis essential for sustaining proliferative and oncogenic signalling in NSCLC. (H) Control and shPEAK1 cells were treated with TGFβ inhibitor and expression of RHOV was measured. The expression of RHOV was increased with TGFβ inhibitor even in presence of shPEAK1 suggesting, PEAK1 mediated inhibition of TGFβ plays a crucial role in RHOV expression.

To validate RHOV’s interaction with PEAK1, we performed pull-down assays, confirming their binding (**Figure 6C**). Investigating PEAK1’s role in RHOV-driven oncogenic signaling, we conducted cell proliferation assays under different conditions. As expected, RHOV overexpression significantly enhanced cell proliferation compared to controls, reinforcing its pro-proliferative role (**Figure 6D**). To assess whether PEAK1 contributes to this effect, we silenced PEAK1 using shRNA, which led to a marked reduction in cell proliferation, indicating that PEAK1 is essential for NSCLC cell proliferation (**Figure 6D**). Furthermore, PEAK1 knockdown in RHOV-overexpressing cells completely abolished RHOV-induced proliferation, demonstrating that PEAK1 is necessary for RHOV-mediated cell growth (**Figure 6D**). To further elucidate the mechanism underlying this interaction, we examined MYC expression under these conditions. As expected, MYC levels were significantly elevated in RHOV- overexpressing cells (**Figures 6E** and **6F**). However, PEAK1 knockdown alone reduced MYC expression, with an even more pronounced effect when PEAK1 was silenced in RHOV- overexpressing cells (**Figures 6E** and **6F**). These findings indicate that PEAK1 plays a crucial role in MYC regulation and that RHOV sustains high MYC levels via PEAK1. Collectively, these results suggest that the RHOV–PEAK1 axis is essential for MYC-driven proliferation in NSCLC, positioning PEAK1 as a key mediator of RHOV’s oncogenic effects. To explore the regulatory mechanisms governing the RHOV-PEAK1-MYC axis, we examined key signaling pathways involved in its modulation. Western blot analysis revealed a significant decrease in phosphorylated AKT and ERK levels in both shRHOV and shPEAK1 samples (**Figure 6G**), suggesting that RHOV and PEAK1 act upstream of AKT and MAPK signaling. This indicates that the RHOV–PEAK1-MYC axis is regulated by the PI3K/MAPK pathway. Interestingly, we observed that PEAK1 knockdown resulted in decreased RHOV expression, suggesting a role for PEAK1 in regulating RHOV levels (**Figure 6F**). Given our previous findings that TGFβ negatively regulates RHOV, we examined TGFβ expression in PEAK1 knockdown cells and found a significant increase (**Figure 6G**). This suggests that PEAK1 suppression is associated with elevated TGFβ expression, which in turn inhibits RHOV. To confirm this, we treated PEAK1 knockdown cells with a TGFβ inhibitor and found that RHOV expression remained unchanged in the presence of the inhibitor (**Figure 6H**).

These findings provide critical insights into the upstream regulation of the RHOV-PEAK1-MYC axis, indicating that RHOV and PEAK1 function within a PI3K/MAPK-dependent pathway. Moreover, PEAK1 regulates RHOV through a TGFβ-dependent feedback loop.

## 5. Discussion

The intricate signaling networks governed by small GTPases are fundamental to cellular function, and their dysregulation is increasingly recognized as a hallmark of cancer. While oncogenic mutations in RAS GTPases are well-established drivers of tumorigenesis, the roles of other GTPase families, particularly the Rho subfamily involved in cytoskeletal dynamics and motility, remain less defined, especially concerning lesser-known members [12,25–27]. This study undertook a systematic investigation into Rho GTPase dysregulation across various cancers, identifying RHOV as a consistently overexpressed gene, particularly in non-small cell lung cancer (NSCLC), a finding corroborated across multiple large-scale datasets (Figure 1A, 1B; Supplementary Figure 1A, 1B). Given the limited prior knowledge regarding RHOV’s function in malignancy, this study aimed to elucidate its biological roles and clinical significance in NSCLC. The findings presented herein establish RHOV as a critical oncogenic driver in NSCLC, demonstrating its overexpression correlates strongly with poor patient prognosis, and revealing its essential functions in promoting cell proliferation and migration through distinct molecular mechanisms involving TGFβ signaling, MYC activation, metabolic reprogramming, and interaction with the PEAK1. Earlier studies have shown the function of RHOV in NSCLC, however, detailed upstream and downstream regulation was lacking [6,28].

A primary finding of this investigation is the consistent and significant overexpression of RHOV in NSCLC tissues compared to corresponding normal lung tissues. This observation was robustly supported by analyses across diverse platforms, including TCGA RNA-sequencing data, multiple independent microarray datasets curated via Oncomine analysis, and was further validated at the protein level using immunohistochemistry data from the Human Protein Atlas (**Figure 1A, 1B, 1D**; **Supplementary Figure 1A, 1B**). The marked difference in expression levels translated into a strong diagnostic potential, as demonstrated by Receiver Operating Characteristic (ROC) analysis, which indicated high specificity and sensitivity for RHOV expression in distinguishing NSCLC tumor samples from normal tissues (**Figure 1C**).

Beyond its diagnostic potential, the clinical relevance of RHOV overexpression was underscored by its strong association with patient outcomes. Analysis of four independent NSCLC patient cohorts, encompassing a total of 1540 patients with associated clinical data, revealed a significant correlation between high RHOV expression and reduced overall survival. Univariate Cox proportional hazards regression analysis across three cohorts with available median survival data consistently identified elevated RHOV levels as a significant predictor of poorer survival (**Figure 1G**). Crucially, multivariate Cox regression analysis, incorporating both RHOV expression and tumor stage, established RHOV as an independent prognostic factor (Figure 1G). This independence suggests that RHOV expression captures biological aspects of tumor aggressiveness beyond those reflected by standard anatomical staging, potentially indicating intrinsic cellular properties driving poor outcomes. Notably, the prognostic significance of RHOV extended to patients diagnosed with early-stage NSCLC (**Supplementary Figure 1B**). Given that prognosis is a critical factor in determining treatment strategies, particularly regarding adjuvant therapies for early-stage disease, RHOV expression emerges as a potentially valuable biomarker to refine risk stratification and guide clinical decision-making in NSCLC management. Corroborating the clinical associations suggesting an oncogenic role, functional studies provided direct evidence that RHOV actively promotes key malignant phenotypes in NSCLC cells. Loss-of-function experiments, employing both transient siRNA-mediated silencing and stable shRNA-mediated knockdown in A549 and H23 lung adenocarcinoma cells, consistently demonstrated that RHOV depletion significantly impairs cell proliferation and clonogenic potential (**Supplementary Figure 2C-H; Figure 2A-H**). Conversely, overexpression experiments in H460 and A549 cells, resulted in accelerated proliferation rates (**Figure 2A-H**). These complementary approaches strongly support a role for RHOV in driving NSCLC cell growth. RHOV knockdown led to a significant reduction in migratory capacity (**Figure 3A-C**; **Supplementary Figure 3A-B**). Mechanistically, this migration defect correlated with distinct alterations in cytoskeletal architecture; phalloidin staining revealed a marked decrease in actin filament-rich protrusive structures, such as lamellipodia or filopodia, at the cell periphery in RHOV-depleted cells compared to controls (**Figure 3D**). These observations were validated using overexpression system (**Figure 3E, F and G; Supplementary Figure 3C-D**). Collectively, these data establish RHOV as a critical regulator of both proliferation and migration in NSCLC cells, exerting its pro-migratory effects, through modulation of actin cytoskeletal dynamics.

Mechanistically, we explored the upstream regulation and downstream effectors of RHOV. We identified a novel regulatory axis involving TGFβ signaling. Analysis of GEO datasets and experimental validation in A549 cells revealed that TGFβ consistently suppresses RHOV expression (**Figure 3G-I**). This regulation appears to be direct and dependent on the canonical SMAD3 pathway (**Figure 3J-L**). Intriguingly, we observed an inverse correlation between RHOV levels and the expression of TGFβ pathway components (TGFβ1, TGFβ2, TGFβR2) in NSCLC patient samples and cell lines (**Figure 3M, N**). This suggests that suppression or loss of TGFβ signaling in NSCLC may contribute to RHOV overexpression, potentially representing a mechanism by which cancer cells escape TGFβ-mediated growth inhibition while retaining pro-migratory signals coordinated by RHOV [29].

To dissect the pathways mediating RHOV’s pro-growth effects, we employed RNA sequencing following RHOV knockdown. This revealed significant alterations in gene expression programs, notably a downregulation of MYC target genes and pathways related to translation, cell cycle, and oxidative phosphorylation (**Figure 4A-C**). We confirmed that RHOV knockdown leads to decreased MYC expression at both mRNA and protein levels (**Figure 5A, B**). Crucially, rescuing MYC expression in RHOV-depleted cells restored cell proliferation (**Figure 5C, D**), indicating that MYC is a critical downstream mediator of RHOV-driven growth. Consistent with the GSEA findings and MYC’s role in metabolism, RHOV knockdown reduced mitochondrial respiration (OCR) and ATP production, effects which were also rescued by MYC overexpression (**Figure 5E-H**). This positions RHOV as a regulator of MYC-dependent metabolic programming essential for NSCLC growth. Thus, RHOV signaling, acting via MYC, not only controls proliferation but also impacts fundamental cellular processes of mitochondrial metabolism and redox homeostasis, creating a state of heightened oxidative stress and DNA damage upon its depletion.

The finding that RHOV knockdown induces ROS-mediated DNA damage suggested that RHOV status might influence cellular sensitivity to chemotherapeutic agents, particularly those targeting DNA integrity or repair. Testing this hypothesis, RHOV knockdown was found to dramatically increase the sensitivity of both A549 and H23 cells to the topoisomerase II inhibitor etoposide (**Figure 5K, L**). In contrast, no significant change in sensitivity was observed for doxorubicin, a DNA intercalator (**Figure 5K, L**; **Supplementary Figure 4D**).

The specific sensitization to etoposide upon RHOV knockdown is mechanistically plausible. Etoposide traps topoisomerase II-DNA complexes, leading to DNA strand breaks. Cells experiencing elevated levels of ROS-induced DNA damage, as observed in RHOV knockdown cells, may be particularly reliant on efficient DNA repair mechanisms, including those involving topoisomerase II. Inhibiting this enzyme under conditions of pre-existing DNA damage likely creates a synergistic cytotoxic effect, leading to increased cell death. Doxorubicin, while also inducing DNA damage, acts primarily through intercalation and generation of free radicals, a mechanism that may not synergize as effectively with the specific metabolic and redox state induced by RHOV depletion. These findings suggest that RHOV expression level could potentially serve as a predictive biomarker for etoposide sensitivity in NSCLC, warranting further investigation for patient stratification or the development of combination therapies targeting RHOV-low tumors or inducing a RHOV-low state.

Seeking direct functional partners, our proteomic analysis of RHOV interactors identified several proteins involved in cytoskeletal regulation and signaling, including PAK1/2 [30], IQGAP1, WASL, and GIT2 (**Figure 5B**), consistent with RHOV’s role in migration. Notably, we identified and validated Pseudopodium Enriched Atypical Kinase 1 (PEAK1) as a key RHOV interactor (**Figure 5C**). PEAK1 itself was found to be essential for NSCLC proliferation (**Figure 5D**). Strikingly, PEAK1 knockdown phenocopied RHOV knockdown by reducing MYC expression and abrogated the enhanced proliferation induced by RHOV overexpression (**Figure 5D-F**). This strongly suggests that RHOV requires interaction with PEAK1 to maintain high MYC levels and drive proliferation. Further investigation indicated that the RHOV-PEAK1 axis operates upstream of the AKT and ERK signaling pathways (**Figure 5G**). We also uncovered a potential feedback loop where PEAK1 knockdown increased TGFβ expression, which in turn suppressed RHOV levels (**Figure 5G**), suggesting a complex interplay regulating this oncogenic axis. The stabilization of RHOV upon TGFβ inhibition in PEAK1 knockdown cells supports this model (**Figure 5H**).

In conclusion, this study identifies RHOV as a novel and clinically significant oncogene in NSCLC. Its consistent overexpression serves as a robust, independent biomarker for poor patient prognosis, potentially aiding in risk stratification, particularly in early-stage disease. Functionally, RHOV drives NSCLC cell proliferation and migration. Mechanistically, RHOV operates through a complex network involving transcriptional repression by TGFβ/SMAD3, essential cooperation with the scaffold protein PEAK1 to sustain MYC expression via AKT/ERK signaling, and consequent modulation of mitochondrial metabolism, ROS homeostasis, and DNA damage pathways. This intricate signaling cascade not only contributes to tumor aggressiveness but also creates a specific therapeutic vulnerability, linking RHOV status to etoposide sensitivity. These findings highlight RHOV and its associated pathways, particularly the RHOV-PEAK1-MYC axis, as promising areas for further investigation and potential targets for novel therapeutic strategies in NSCLC.

## Acknowledgements

Sudhanshu Shukla would like to acknowledge Science and Engineering Research Board (SERB), Govt. of India (Grant #ECR/2018/000528) Indian Council for medical research, Govt. of India (Grant # 2021-9513/CMB/ADHOC-BMS and EMDR/SG/13/2023-0244), Department of Biotechnology, Govt of India (Grant id – BT/PR51308/MED/30/2511/2023) and Anusandhan National Research foundation, Govt. Of India (Grant ID – CRG/2023/000837 for the funding. LLM based AI databases were used for the paraphrasing and Grammer correction.

## Author contributions

AC: Conceptualization, Methodology, Formal analysis, Investigation, review and editing.
DA: Investigation
NB: Investigation
PB: Investigation
BKC: Review and editing, Supervision
SS: Conceptualization, Methodology, Formal analysis, Writing – Review & Editing, Funding Acquisition.

