## Supplementary figure for "TGF/J Regulated Small GTPase RHOV interact with PEAK1 and drive MYC Expression to Promote Cellular Proliferation, Migration and Etoposide resistance"

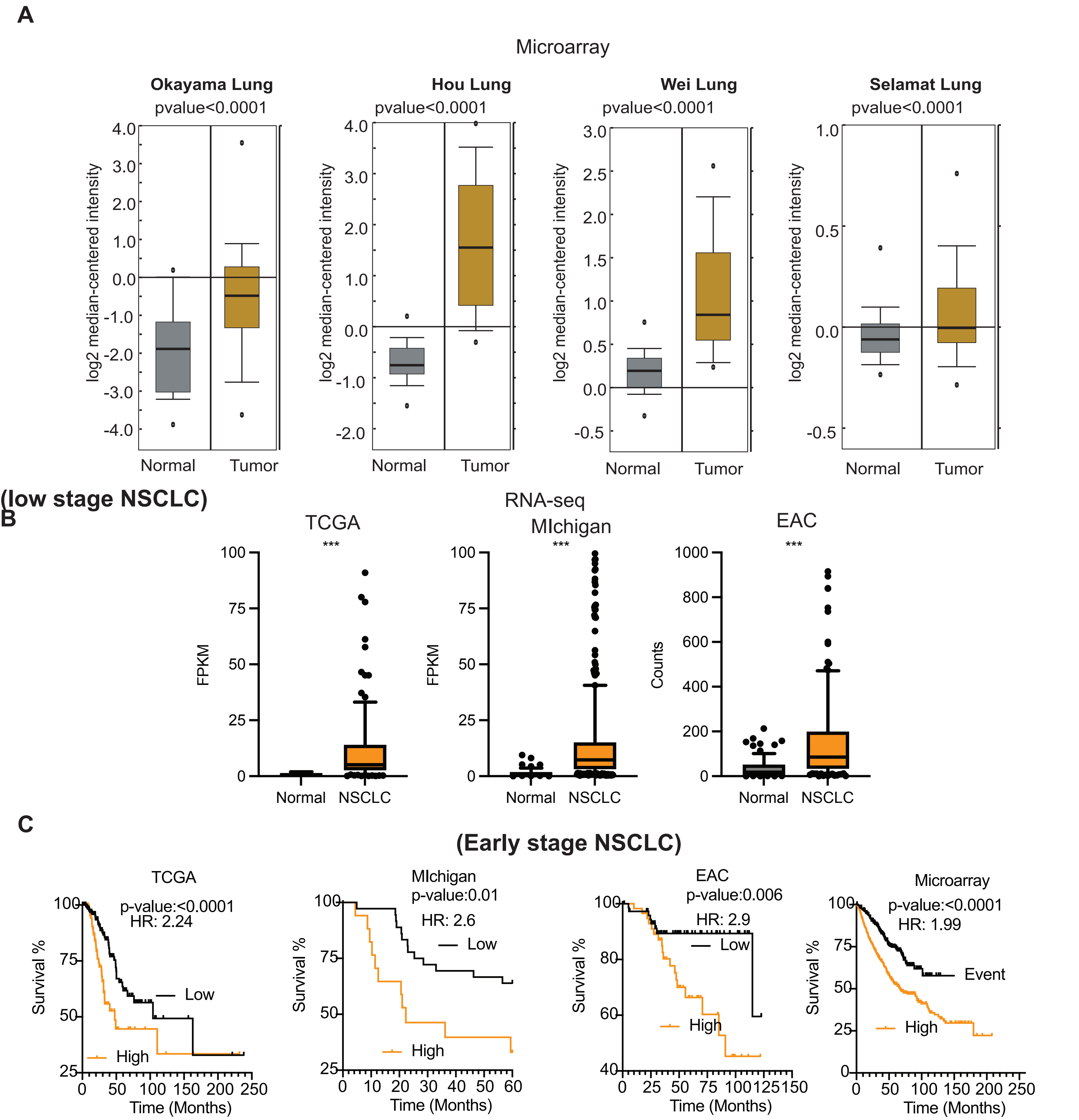


**Supplementary figure 1. (A)** Expression of RHOV was measured in microarray data sets using Oncomine. All the datasets show significant increase in expression RHOV in NSCLC samples than in Normal.

**(B)** Expression of RHOV was measured in RNA-seq datasets. All the three datasets show significant increase in expression RHOV in NSCLC samples than in Normal.

**(C)** Kaplan–Meier survival analyses correlating RHOV expression with overall survival in Early stage NSCLC across TCGA, microarray, Michigan and EAC cohorts. In All the data sets, high RHOV expression is associated with significantly reduced survival (TCGA: HR = 2.24; P <0.0001). (Michigan; HR = 2.26; P = 0.01), ( EAC: HR = 2.9; P = 0.006) and

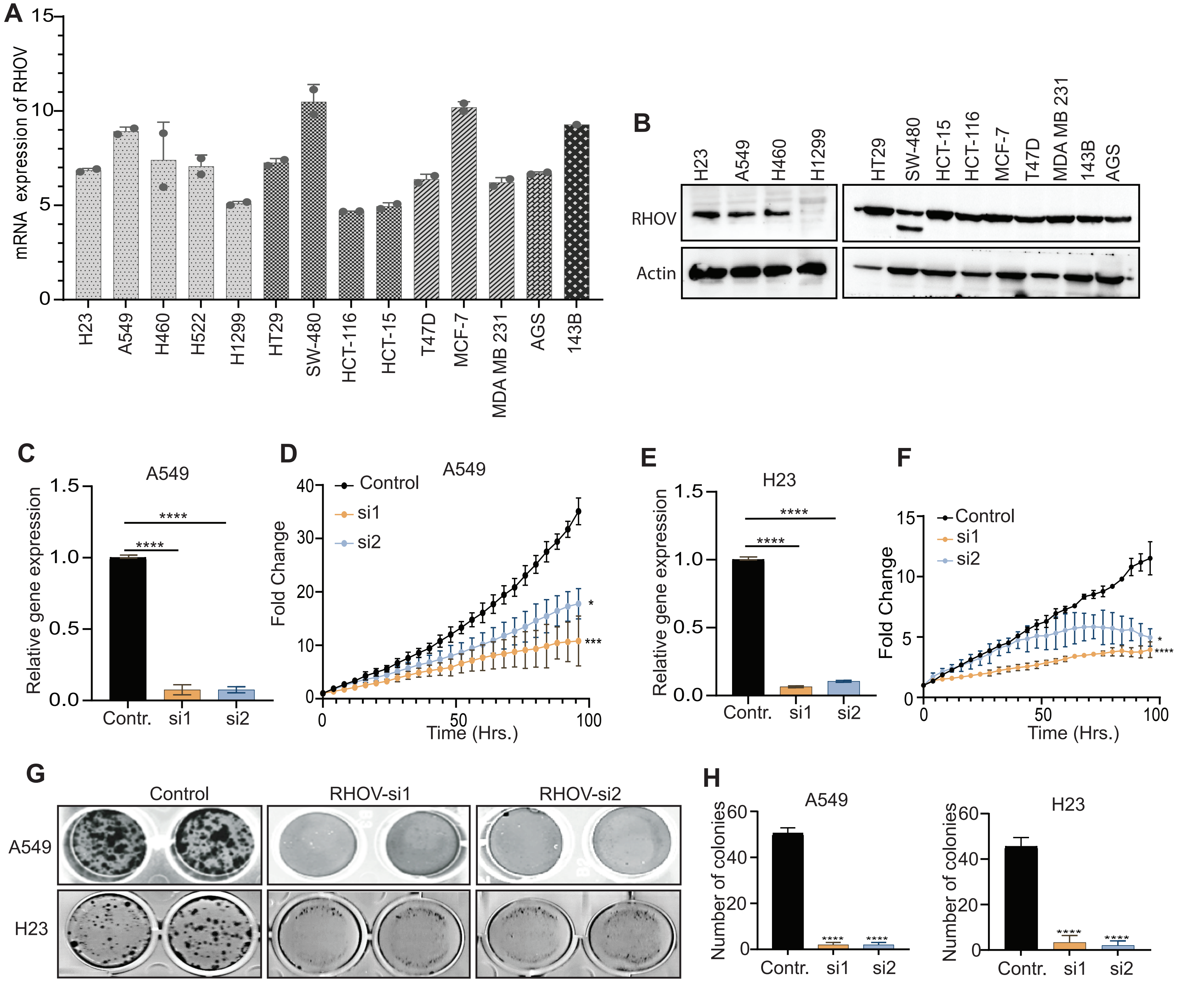


**Supplementary figure 2: (A)** Expression of RHOV in different cell lines were measured using qRT-PCR. dCT values calculated using TPT control are plotted.

**(B)** Expression of RHOVin  different cell lines were measured using western blot.

**(C)** Expressio  of RHOV was measured using qRT-PCR in control A549 and siRHOV A549 cells.

**(D)** Proliferation of control and siRHOV A549 cells were measured using Incucyte. Fold change in confluency is plotted.

**(E)** Expressio  of RHOV was measured using qRT-PCR in control H23and siRHOV H23 cells.

**F)** Proliferation of control and siRHOV H23 cells were measured using Incucyte. Fold change in confluency is plotted.

**(G, H)** Colony supression assay was performed in control and siRHOV cells. After Incubation, cells were stained and colonies were counted and plotted (H).

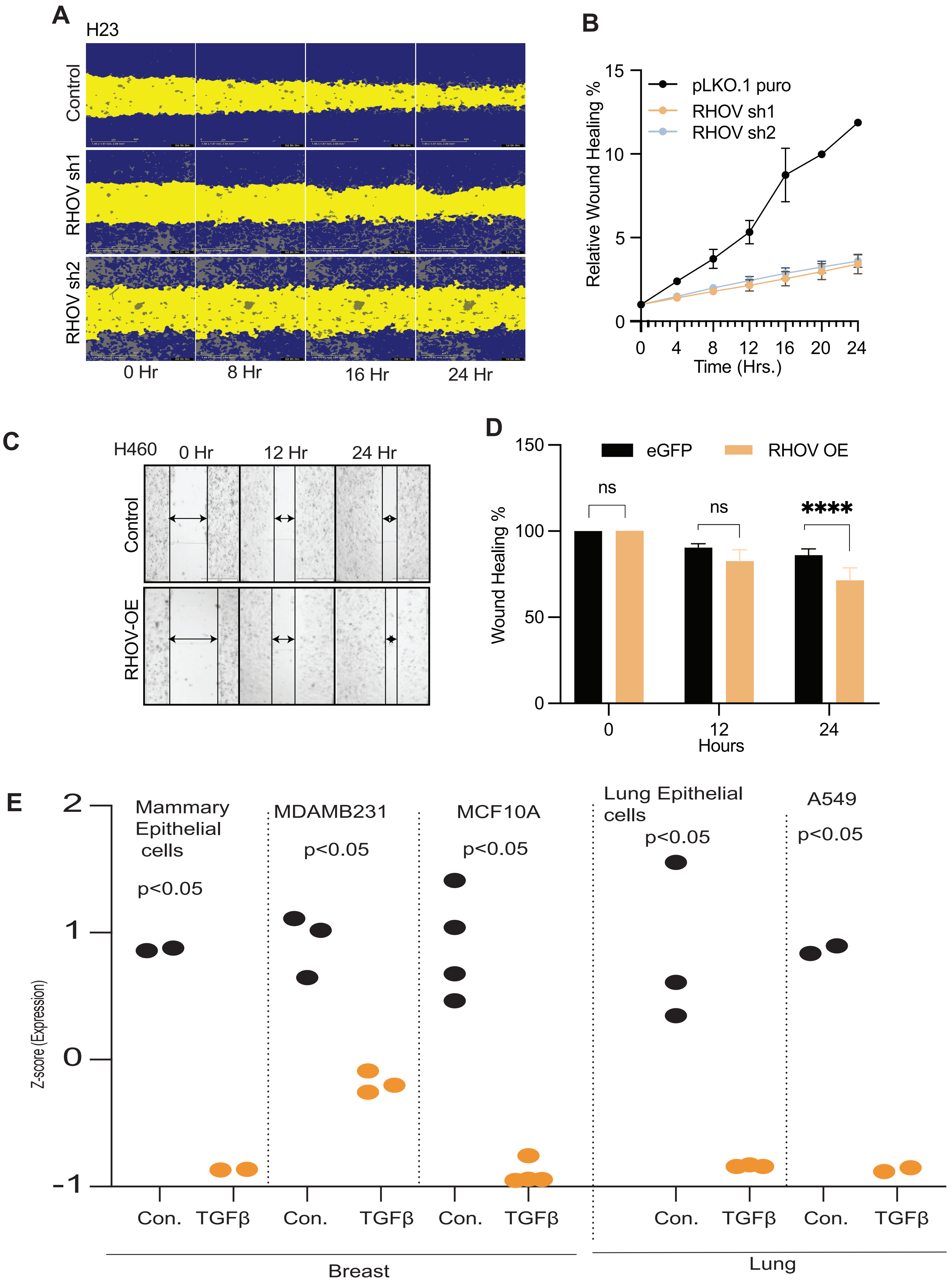


**Supplementary Figure 3: (A)** Control and shRHOV expressing H23 cells were plated and scratch was created. The wound healing was measured using Incucyte at different time points.
**(B)** The wound helaing of Control and shRHOV expressing H23 cells were measured using Incucyte and plotted.
**(C)** Control and RHOV over expressing H460 cells were plated and scratch was created. The wound healing was measured using microscope at different time points.
**(D)** The wound helaing of Control and RHOV over expressing H460 cells cells were measured using microscopy and plotted.
**(E)** Expression of RHOV was measured in differnt menioned cells (data was obtained from GEO) and plotted. Each cell was treated with control and TGFβ. The RHOV expression was reduced after TGFβ treatment.

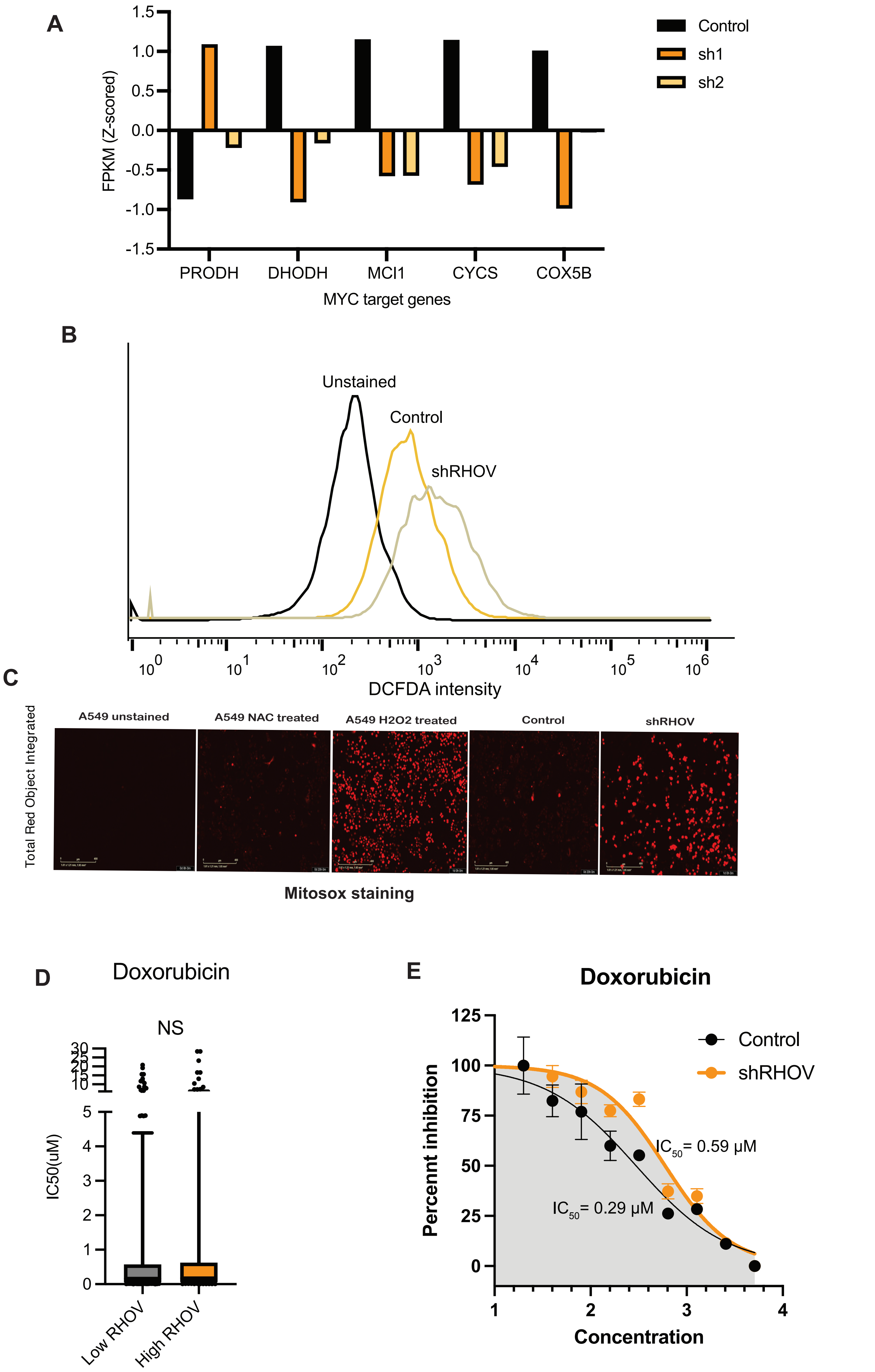


Supplementary figure 4: **(A)** The expression of differnt MYC targets (PRODH-Negative, others-positive) genes invloved in electron transport chain was measured from RNA-seq data of control and shRHOV cell and plotted.
**(B)** A549 control and shRHOV cells were treated with DCFDA and flow cytometery was used to measure the intensity.
**(C)** A549 cells were treated with NAC and H2O2 and stained with Mitosox to measure the mitochondrial ROS. Similarly, mitochondrial ROS was measured in control and shRHOV cells.
**(D)** The Doxorubicin sensitivity of cells lines with high and low expression of RHOV was compared from Genomics of Drug Sensitivity in Cancer database. RHOV high expressing cells showed no significant differnce in IC50 than RHOV low expressing cells (p-value=NS Mann-Whitney test).
**(E)** Doxorubicine induces cytotoxic effect inducing DNA damage. The RHOV knockdown cells show no significant differnce in doxorubicine sensitivity in A549 cells.

A

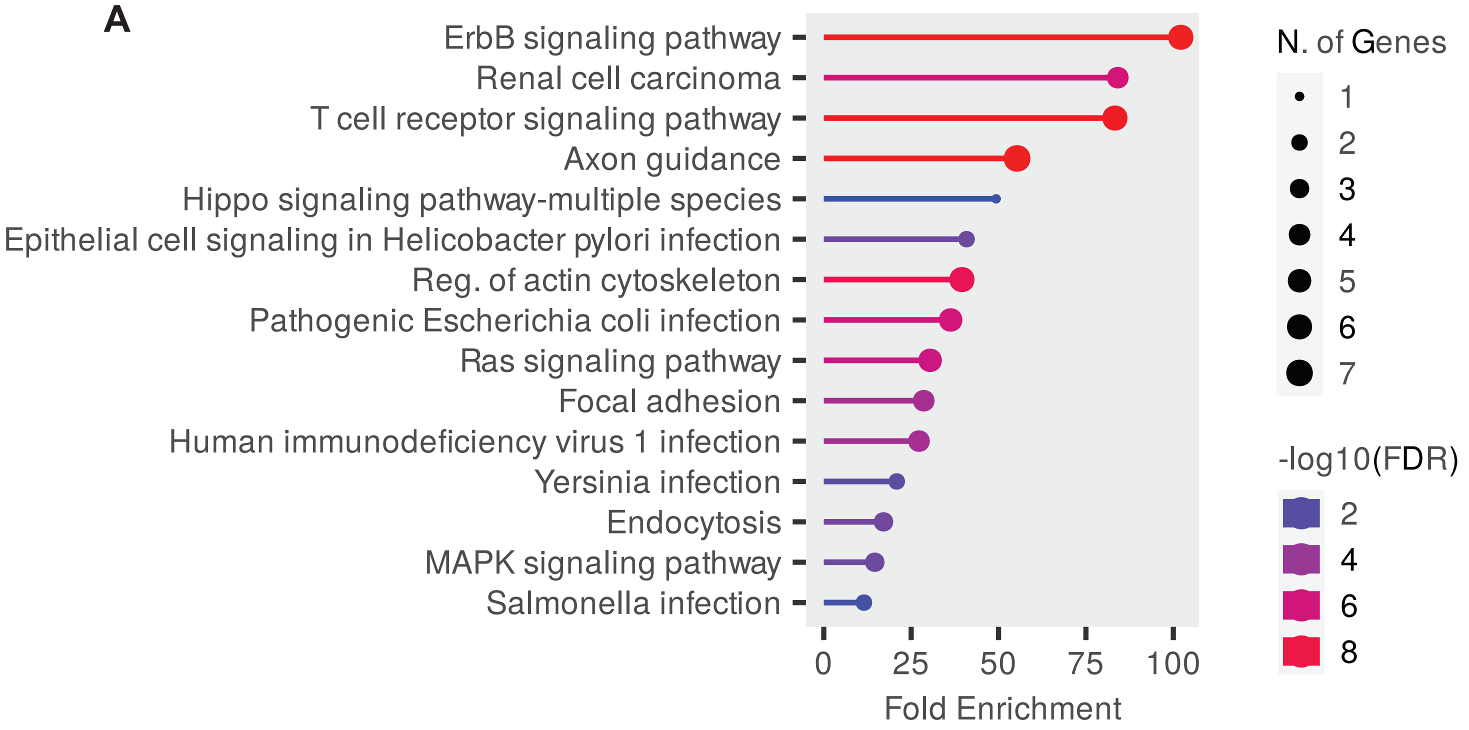


B

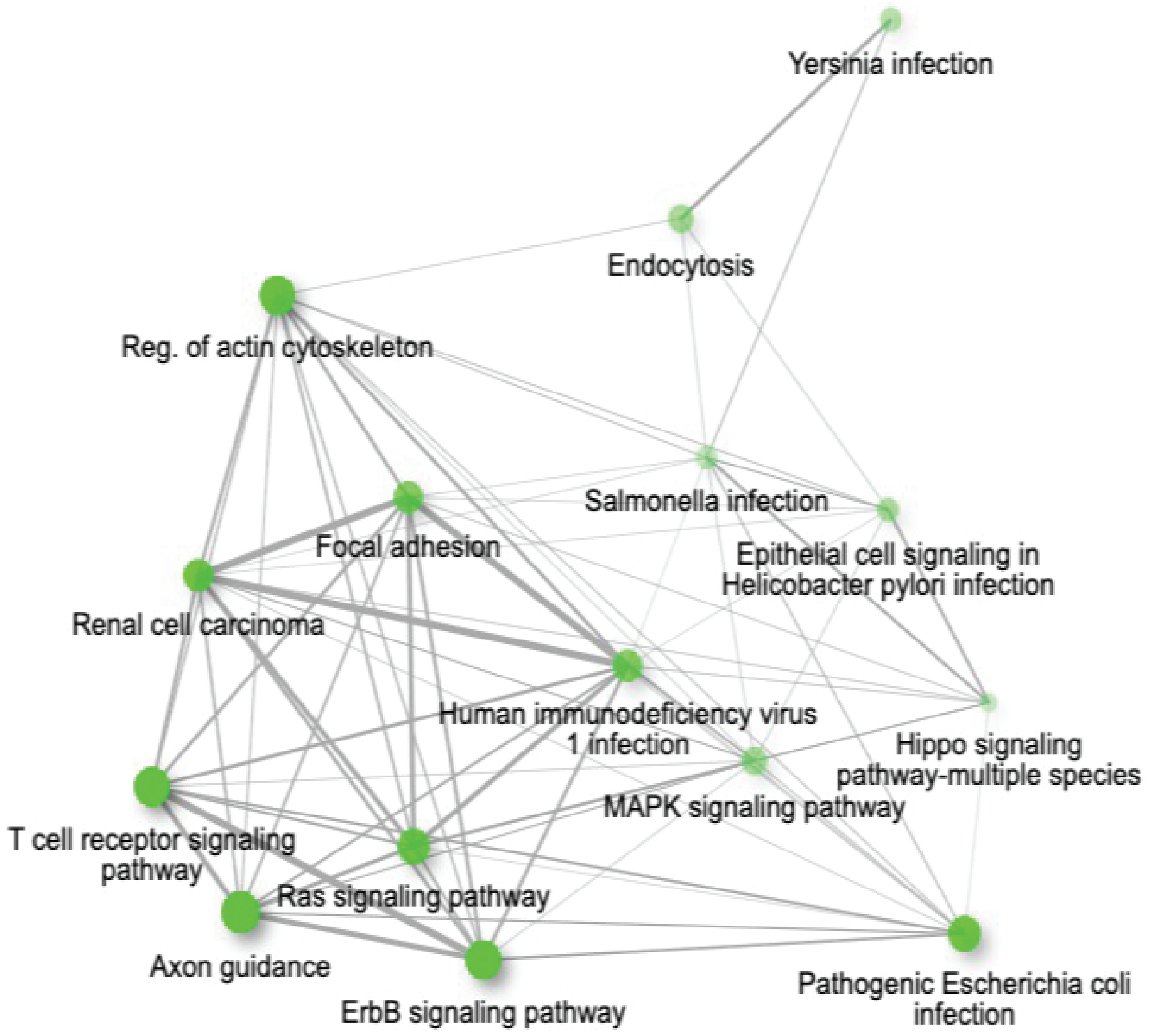


**Supplementary Figure 5: A and B:** The patway analysis of RHOV interacting genes was done using ShineyGo (https://bioinformatics.sdstate.edu/go/) and enriched pathway are plotted. (B) The network of enriched pathways are shown.
